# Single-cell transcriptomic profiling of human pancreatic islets reveals genes responsive to glucose exposure over 24 hours

**DOI:** 10.1101/2023.06.06.543931

**Authors:** Caleb M. Grenko, Lori L. Bonnycastle, Henry J. Taylor, Tingfen Yan, Amy J. Swift, Catherine C. Robertson, Narisu Narisu, Michael R. Erdos, Francis S. Collins, D. Leland Taylor

## Abstract

Disruption of pancreatic islet function and glucose homeostasis can lead to the development of sustained hyperglycemia, beta cell glucotoxicity, and ultimately type 2 diabetes (T2D). In this study, we sought to explore the effects of hyperglycemia on human pancreatic islet (HPI) gene expression by exposing HPIs from two donors to low (2.8mM) and high (15.0mM) glucose concentrations over 24 hours, assaying the transcriptome at seven time points using single-cell RNA sequencing (scRNA-seq). We modeled time as both a discrete and continuous variable to determine momentary and longitudinal changes in transcription associated with islet time in culture or glucose exposure. Across all cell types, we identified 1,528 genes associated with time, 1,185 genes associated with glucose exposure, and 845 genes associated with interaction effects between time and glucose. We clustered differentially expressed genes across cell types and found 347 modules of genes with similar expression patterns across time and glucose conditions, including two beta cell modules enriched in genes associated with T2D. Finally, by integrating genomic features from this study and genetic summary statistics for T2D and related traits, we nominate 363 candidate effector genes that may underlie genetic associations for T2D and related traits.

## Introduction

Type 2 diabetes and related complications are among the leading causes of death globally (1). Clinical and genetic studies highlight the central role of pancreatic islet dysfunction and disrupted glucose homeostasis in the development of sustained hyperglycemia and type 2 diabetes (reviewed in (2,3)). Within the pancreatic islet, multiple cell types have been implicated in type 2 diabetes progression, most notably beta cells that secrete insulin in response to glucose stimulation (4), but also other cell types including alpha cells (5) and delta cells (6). Common variant genetic association studies have identified >400 distinct genetic signals associated with type 2 diabetes and type 2 diabetes-related traits (7,8), promising to deliver clues into the genes and molecular pathways that underlie type 2 diabetes development and progression. However, most of the genetic signals identified lie outside protein-coding genes, masking the “effector genes” responsible for driving the genetic association.

One approach to help identify and understand the genes that contribute to type 2 diabetes pathogenesis and progression is to explore the effects of physiologically relevant conditions, such as hyperglycemia, on human islet gene expression. To date, human islet glucose-stimulus studies have shown the effects of hyperglycemia on genes related to insulin secretion (9) and oxidative stress (10,11). The molecular picture from these studies is consistent with our understanding of type 2 diabetes pathophysiology; under normal conditions, transient increases in blood glucose stimulate insulin secretion. However, under the sustained hyperglycemic conditions that occur in type 2 diabetes, the continual demands of insulin production lead to glucotoxicity and apoptosis of beta cells (12) (and possibly other islet cell types (13))—further exacerbating the type 2 diabetes condition. Despite the known importance of multiple islet cell types in the context of diabetes (4–6), to date, transcriptomic studies on the effects of hyperglycemia on primary human islets have used gene expression readouts from bulk islet tissue (9–11), thus masking cell-type specific expression signatures which may be relevant to diabetes.

In this study, we perform single-cell RNA sequencing (scRNA-seq) to measure the transcriptional changes associated with sustained glucose exposure across islet cell types. We expose human pancreatic islets from two donors to euglycemic conditions (2.8mM) and hyperglycemic conditions (15.0mM) *in vitro* and sample cells at seven time points over 24 hours. We identify genes with changes in expression associated with time in culture or glucose exposure using both discrete models, which test for changes in expression at a given time point, and continuous models, which analyze all of the data longitudinally across time points. Using these models, we test for three types of gene expression effects: (i) time in culture, (ii) glucose exposure, and (iii) interaction effects between time and glucose exposure (time:glucose; i.e., how gene expression patterns change between low and high glucose through time). Through this time-series experimental design, we demonstrate the importance of controlling for “time in culture” to isolate expression effects associated with glucose exposure. We show that the transcriptomic response associated with high glucose exposure (total number of associated genes) plateaus at 8 hours in beta cells and is variable across other cell types. For beta cells specifically, we find multifactorial effects on gene expression whereby many genes have patterns associated with all three of the considered variables: time, glucose exposure, and time:glucose—highlighting the sensitivity of beta cells to both time in culture and glucose exposure. By clustering the expression profiles of genes in euglycemic and hyperglycemic conditions through time, we identify cell-type specific modules of co-expressed genes. For two of the beta cell expression modules, we find enrichment in genes related to type 2 diabetes, one of which shows strong enrichment in pathways related to insulin secretion. Finally, by modeling genetic association signals from type 2 diabetes and type 2 diabetes-related traits using the data from this study, we identify 363 candidate effector genes that may drive genetic association signals for type 2 diabetes, blood glucose, and glycated hemoglobin (HbA1c)—seven of which feature in the aforementioned type 2 diabetes-associated expression modules. Collectively, the results from this study provide a high-resolution view of the effects of euglycemic and hyperglycemic conditions on islet cell types through time and should help guide future experiments to understand the molecular mechanisms that lead to islet dysfunction in disease states like type 2 diabetes.

## Results

### Single-cell RNA sequencing of glucose-stimulated human pancreatic islets

We obtained human pancreatic islets from two donors and acclimated the islets to a basal, euglycemic media of 2.8mM glucose (50mg/dL) over one hour (Fig. 1). At the end of one hour, we sampled a subset of cells and performed scRNA-seq (Methods). For the remaining cells, we either maintained them in the low (basal) glucose concentration or exposed them to a high (hyperglycemic) glucose concentration of 15.0mM (270mg/dL). We subsequently sampled cells at six time points for scRNA-seq over the course of 24 hours (cells remained in the glucose concentration over the entire time course). We refer to cells from islets sampled after one hour of acclimation in 2.8mM glucose as “basal” cells (time point 0 hours) and cells from islets sampled at later time points as either “low” or “high” glucose cells (time point 1-24 hours). After quality control procedures (Methods), we obtained 49,895 cells spanning eight cell types, including endocrine cells (alpha 30%, beta 29%, delta 5.2%, and gamma 3.4%), exocrine cells (acinar 20% and ductal 12%), macrophages (0.42%), and endothelial cells (0.15%; Fig. S1-S3). For subsequent analysis, we removed macrophage and endothelial cells as these cells were rare and poorly represented across donors, time points, and glucose conditions (<80 cells at each time point; Fig. S3).

**Fig. 1.**
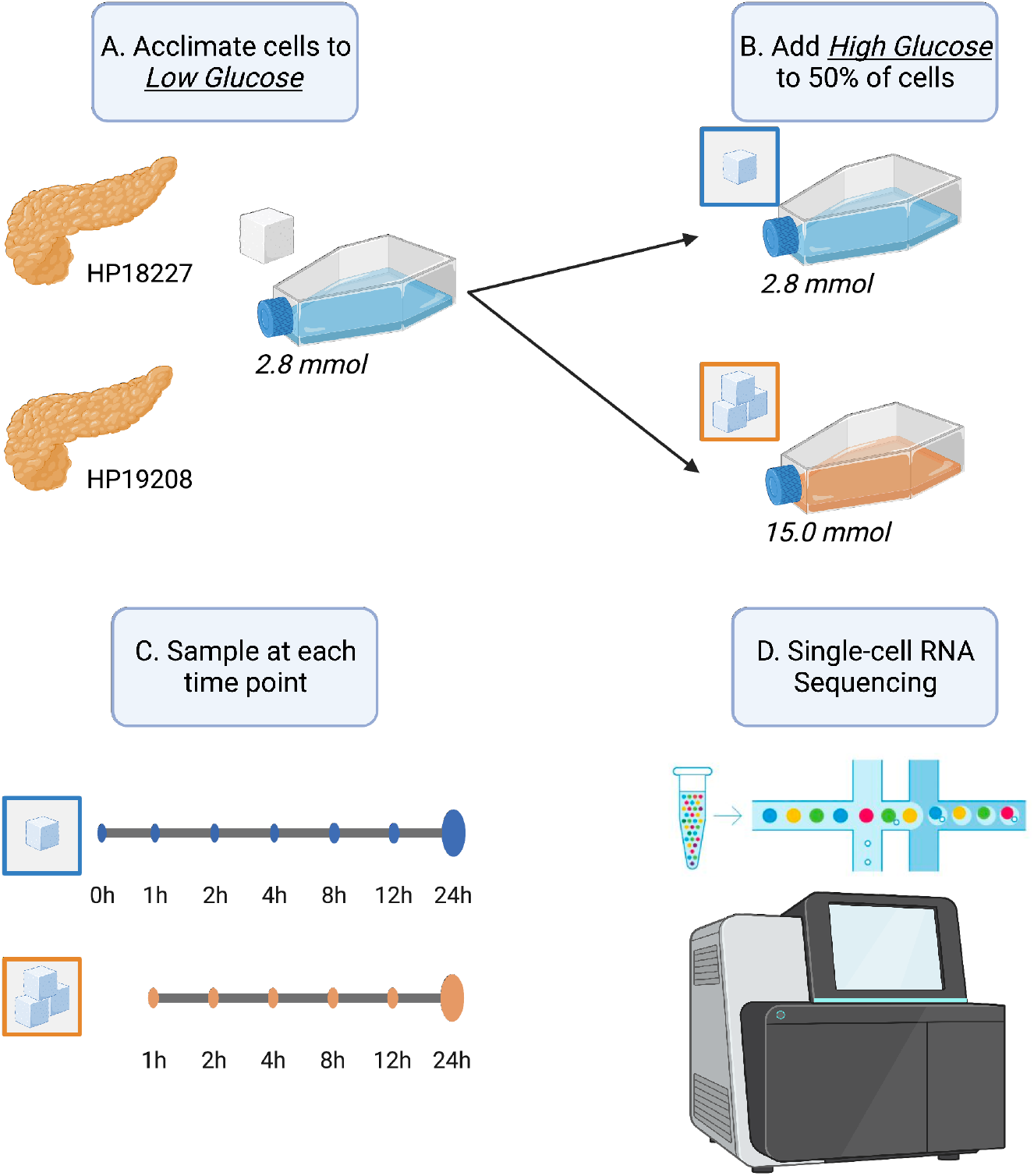
Graphical overview of this study. (A) Upon receipt, we acclimated pancreatic islets from two donors to low glucose conditions (2.8 mmol) for one hour. (B) After acclimation, we exposed half of the islets to high glucose (15.0 mmol) and kept the other half in low glucose conditions (2.8 mmol). (C) At 0, 1, 2, 4, 8, 12, and 24 hour time points, we sampled islets and (D) performed single-cell RNA sequencing. Time point 0 corresponds to cells after one hour low glucose acclimation, prior to starting the stimulation experiment.

### Time point specific effects of glucose induction

We fit three discrete models to characterize the transcriptomic response of islet cell types to glucose stimulation at each time point (Methods). First, in the basal versus high glucose model (“basal-versus-high” or “BvH”), we compared the gene expression of cells in the basal state (2.8mM glucose, 0 hour time point) to cells in high glucose at each time point. This model identifies transcriptomic effects of high glucose at various time points but confounds the impact of high glucose with time in culture. Second, in the basal versus low glucose model (“basal-versus-low” or “BvL”), we compared the expression of cells in the basal state (2.8mM glucose, 0 hour time point) to cells that remained in the low glucose condition at every other time point—isolating time in culture effects while removing the glucose concentration as a confounding factor. Finally, we fit a third model within each time point, comparing cells exposed to low glucose against cells exposed to high glucose (“low-versus-high” or “LvH”) to identify glucose-related effects while controlling for time.

For the BvH and BvL models, the number of associated genes increased over time across all cell types (Fig. 2A). By contrast, for the LvH model (which best isolates glucose-related effects), the total number of associated genes peaked or plateaued between 8 and 12 hours depending on the cell type (Fig. 2A). For all cell types other than beta cells, the proportion of genes differentially expressed in both the BvH and BvL models increased across time, suggesting that time in culture may confound the BvH results, as anticipated (Fig. 2B). Indeed, when we compared the signed -log_10_(*P*-values) across models, we found that the BvH and BvL values were strongly correlated, but not the LvH signed -log_10_(*P*-values) (Fig. S4). Combined, these results confirm that (i) the BvH model confounds tissue and glucose effects and (ii) the BvL and LvH models best isolate time and glucose effects, respectively. Therefore, we chose to focus on the BvL and LvH models for subsequent analyses.

**Fig. 2.**
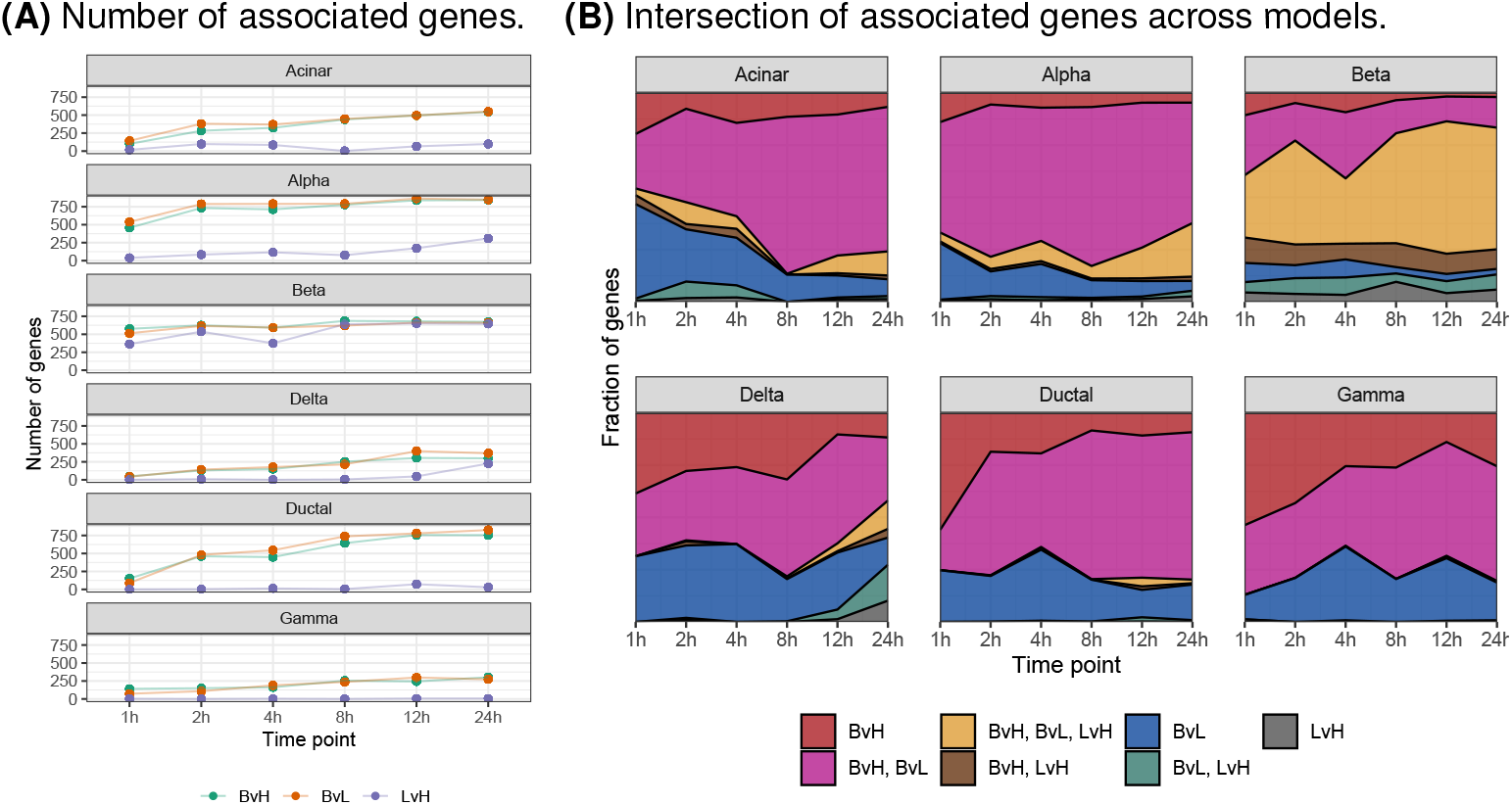
Differentially expressed genes in discrete time models. (A) Number of associated genes (FDR*<*5%; y-axis) at each time point (x-axis) for three models that treat time as a discrete variable: basal-versus-low (BvL, green), basal-versus-high (BvH, orange), and low-versus-high (LvH, purple). (B) Fraction of associated genes (y-axis) that are model-specific or shared across models at each time point (x-axis). Color denotes model or combination of models.

To further characterize the transcriptional response to time in culture and glucose exposure, we calculated (i) the overlap of associated genes (FDR<5%) across time points within each model and cell type (Fig. S5-S6) and (ii) the earliest time point of when a gene showed an association (Fig. S7). Focusing on the LvH model (Fig. S6), the time point with the greatest number of uniquely associated genes varied across cell types: 24 hours for alpha, delta, and gamma cells, 12 hours for ductal cells, 8 hours for beta cells, and 2 hours for acinar cells. Notably, for beta cells, the number of associated genes shared across all time points greatly exceeded the number of genes exclusively expressed at any time point—indicating a rapid and sustained transcriptomic response of beta cells to glucose exposure. The rapid transcriptomic response of beta cells to glucose exposure was also apparent when we considered the earliest time at which a differentially expressed gene was identified; in beta cells, roughly 75% (1,721/2,310) of all genes differentially expressed in the LvH model at 4 hours or later already showed expression differences at 1-2 hours (Fig. S7). This trend was unique to beta cells.

### Temporal dynamics of gene expression in response to glucose induction

While discrete models can effectively identify time-specific effects (Fig. S5-S6), such models fail to make use of the full potential of the data as they do not simultaneously model gene expression across all time points, leading to (i) reduced power to detect effects common to multiple time points and (ii) an inability to model more complex relationships like interaction effects between time and glucose concentration (time:glucose effects). Treating time as a continuous variable, we fit a series of models to identify gene expression patterns associated with (i) time (with glucose as a covariate), (ii) glucose concentration (with time as a covariate), and (iii) time:glucose (with time and glucose as covariates; Methods).

To model time, we considered two metrics: (i) sampled time (i.e., the experimental time point) and (ii) interpolated time (Fig. S8-S9), a metric that models similarities between cells based on the assumption that cells sampled at various time points are not synchronized at the exact same response phase (i.e., some cells sampled at 8 hours may have lagged in their response to stimulation and therefore have an expression profile more similar to 6 hours than 8; Methods). We compared the results of time, glucose, and time:glucose models fit using the two different time metrics and found a slight boost in power using interpolated time over sampled time with concordant directions of effect (Fig. S9B-C), suggesting that interpolated time more accurately represents the phase of cellular response than sampled time. Therefore, in all subsequent continuous models, we used interpolated time.

Across the three continuous models, we identified 1,431 genes with expression patterns associated with time, glucose, and time:glucose (Fig. S9C). As anticipated, compared to the discrete models, the continuous models identified many effects that were not detected previously (Fig. S10).

We calculated the overlap of associated genes between models within cell types (Fig. 3A) and between cell types within models (Fig. 3B). We found that across the three models within a given cell type, very few genes exhibited exclusive glucose effects (Fig. 3A). By contrast, a large number of genes were associated exclusively with time in culture, a trend particularly strong in ductal and gamma cells but absent in beta cells. In beta cells, most genes showed expression changes associated with time, glucose, and time:glucose interaction effects. These included several type 2 diabetes-related genes, such as *ABCC8* (14,15), *SLC30A8* (16,17), *PCSK1* (18–21), *PAX6* (22–24), and *G6PC2* (25–28). When we considered the overlap of genes across cell types within each model (Fig. 3B), we found that the genes associated with time in culture showed dispersed trends: 8.7% (124/1422) were shared across all cell types, 57% (810/1422) were shared between a grouping of cell types, and 34% (488/1422) were cell type specific. For genes associated with glucose and glucose:time, genes tended to be cell type specific (59%, 527/890 for glucose; 57%, 487/845 for glucose:time).

**Fig. 3.**
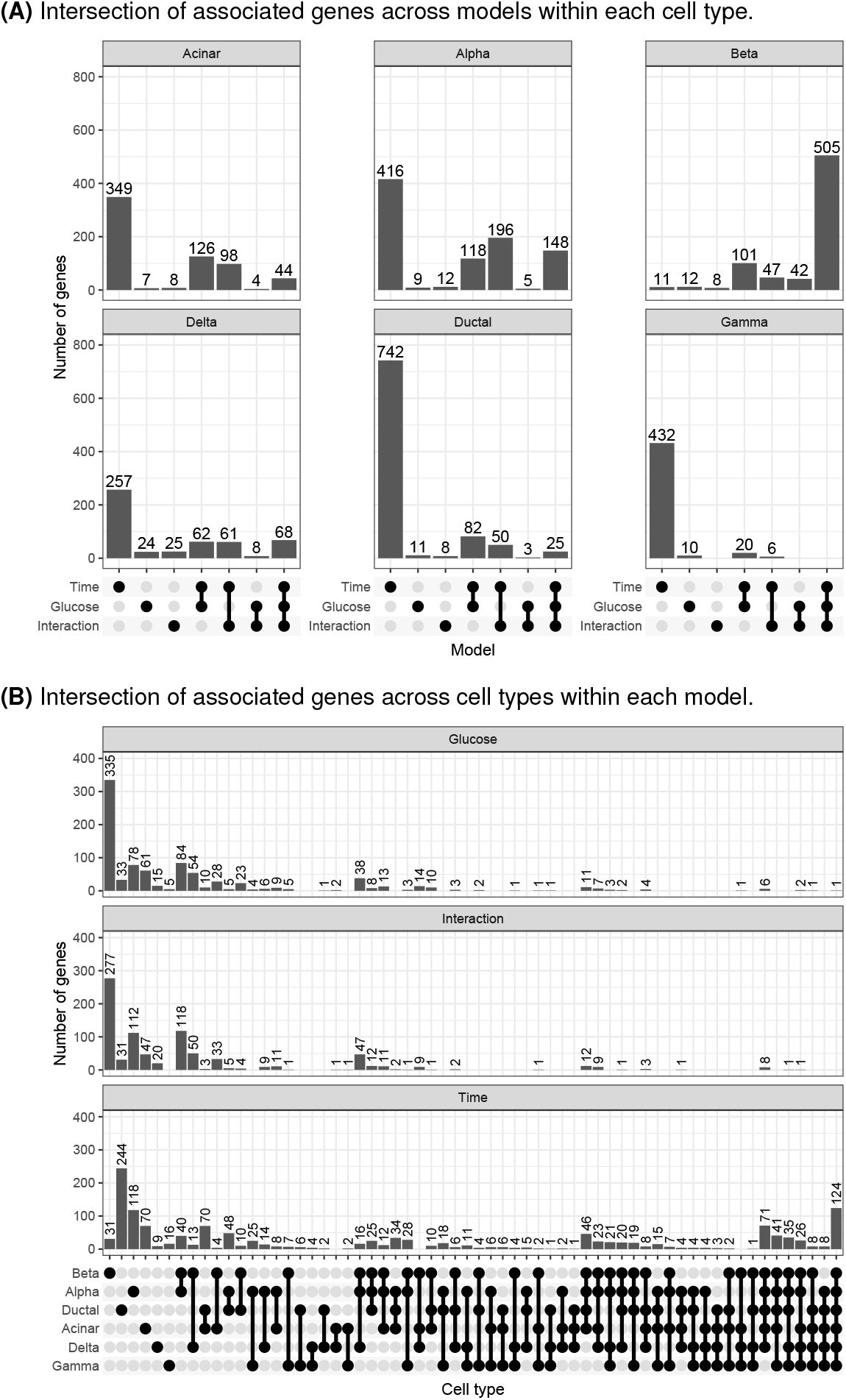
Differentially expressed genes in continuous time models. (A) Number of associated genes (FDR*<*5%; y-axis) in each cell type that are model-specific or shared across models (x-axis). (B) Number of associated genes (FDR*<*5%; y-axis) from each model that are cell type-specific or shared across cell types (x-axis).

### Identification of type 2 diabetes-relevant gene expression modules across time

We selected genes differentially expressed in any of the models, excluding the BvH model, and clustered their expression patterns over time within each cell type to learn modules of co-expressed genes (Methods). To ensure that the final modules were stable, we tested different clustering parameter configurations and generated the final cluster modules using the frequency in which genes co-clustered together, requiring that all genes in the final modules co-cluster ≥75% of the time and that each module contain ≥5 genes. In total, we identified 347 modules across all cell types (Table S2).

We tested each module for enrichment in genes annotated as a type 2 diabetes effector gene in the Type 2 Diabetes Knowledge Portal (https://t2d.hugeamp.org; Methods). We identified two enriched expression modules (FDR<5%, hypergeometric test), both in beta cells: modules 84 and 103 (Fig. 4A-B). Module 84 contained three effector genes from the Type 2 Diabetes Knowledge Portal—*ABCC8, SLC30A8*, and *G6PC2—*and module 103 included two effector genes: *PAX6* and *STARD10*. To better understand the function of these modules, we calculated the enrichment of gene ontology (GO) terms within each module (Methods). Module 84 was enriched in terms related to insulin secretion and module 103 was enriched in terms related to RNA binding and regulation (FDR<5%; Fig. 4C,S11).

**Fig. 4.**
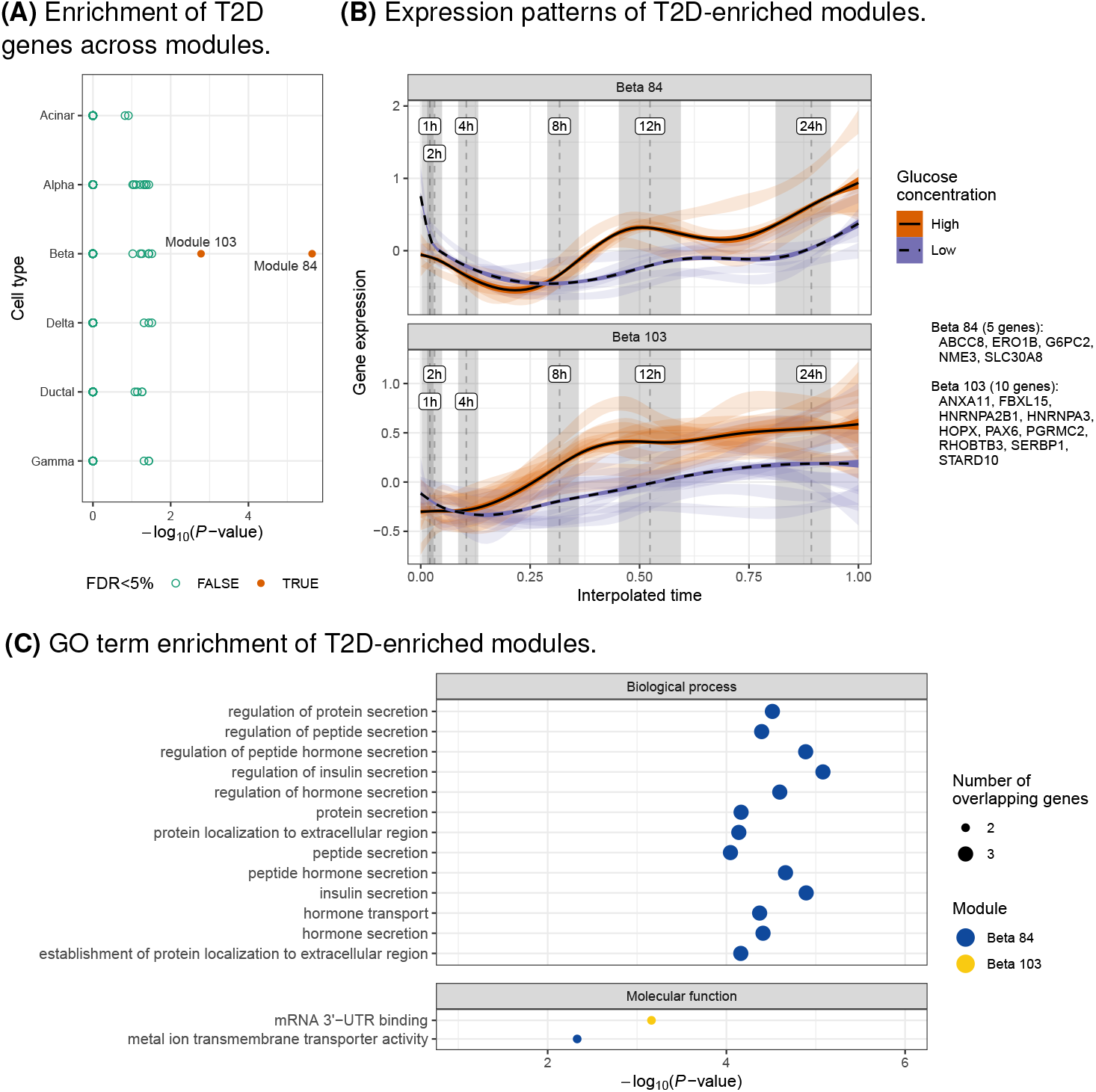
Characterization of gene expression modules related to T2D. (A) Enrichment (x-axis; hypergeometric test) of T2D genes (https://t2d.hugeamp.org) in gene expression modules across cell types (y-axis). (B) Expression of genes (y-axis; CP10K z-scores calculated within each gene) in modules enriched for T2D genes (FDR*<*5%) across time (x-axis) and glucose concentrations (color). Light ribbons correspond to individual genes, where the width depicts the 95% confidence interval of the regression that summarizes gene expression. Dark lines and ribbons show the regression across all genes. Dashed vertical lines show the median interpolated time mapping to standard time and gray shaded regions denote the interquartile range. Genes in each module are listed in the figure guide. (C) Enrichment (x-axis) of GO terms (y-axis) for modules enriched for T2D genes (FDR*<*5%; color), faceted by ontology class. Point size corresponds to the number of overlapping genes. Only terms with FDR¡5% shown and 2 genes overlapping expression modules.

### Nomination of candidate effector genes for type 2 diabetes and type 2 diabetes-related traits

In addition to prioritizing genes that may be related to type 2 diabetes through co-expression patterns, we sought to identify additional candidate effector genes for type 2 diabetes and type 2 diabetes-related traits by modeling genetic association summary statistics directly. Using the Polygenic Priority Score (PoPS) method (29), we modeled genetic association summary statistics for type 2 diabetes (7), blood glucose (Sinnott-Armstrong et al., 2021), and HbA1c (Sinnott-Armstrong et al., 2021) using genomic features derived from this study (e.g., cell type expression specificity patterns, differential expression test statistics; Methods).

We identified 363 unique candidate effector genes across all three phenotypes (FDR<5%): 14 for type 2 diabetes, 358 for blood glucose, and 3 for HbA1c (Fig. 5). We compared the -log_10_(*P*-value) for each gene across phenotypes and found them to be slightly correlated (minimum Pearson’s *r*=0.3), indicating the presence of both common and disease/trait-specific effects (Fig. S12). Among the candidate effector genes, 351 (97%) were associated with only one phenotype and 12 (3.3%) were associated with two phenotypes. Of the 132 type 2 diabetes Knowledge Portal effector genes, we identified 18 (14%) as candidate effector genes for at least one phenotype, including *ABCC8, KCNJ11, NKX2-2, G6PC2, PAM, HNF4A, SLC30A8, RFX6, PCSK1*, and *GLIS3*.

**Fig. 5.**
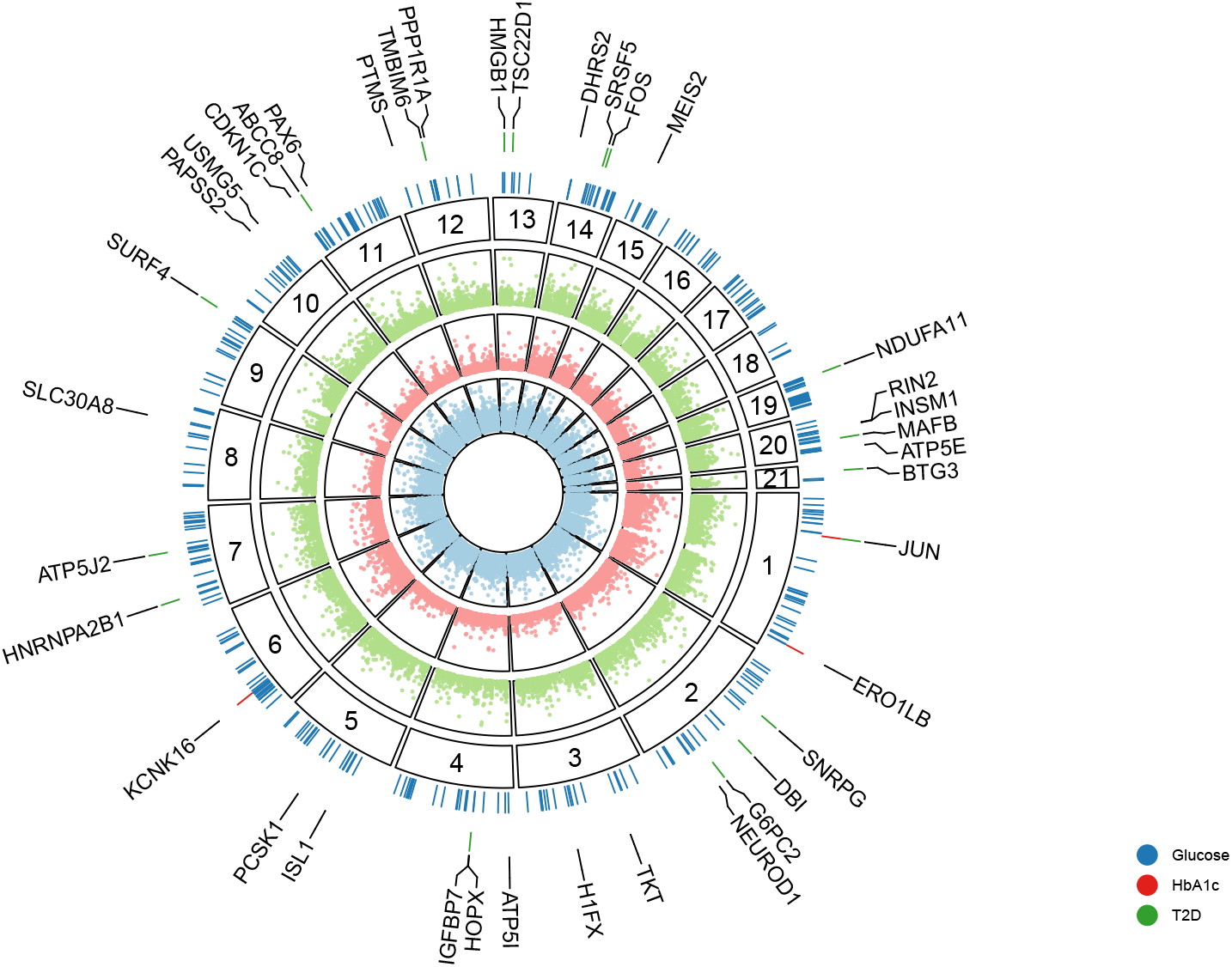
Candidate effector genes. Empirical -log_10_(*P*-values) for each gene (points) prioritized by PoPS analysis using genetic association summary statistics for blood glucose (blue), HbA1c (red), and T2D (green). Segments correspond to the chromosome location of genes (numerical labels). Colored ticks in the outer layer depict candidate effector genes across phenotypes (FDR*<*5%). Labeled genes are the T2D, HbA1c, and 25 blood glucose genes with the smallest *P*-values.

We sought to add further evidence for a type 2 diabetes-relevant role of the genes within the type 2 diabetes-enriched beta cell expression modules and intersected the genes in modules 84 and 103 with the 363 candidate effector genes. Eight of the 15 genes in these modules (53%) were identified as candidate effector genes, including four genes not in the Type 2 Diabetes Knowledge Portal effector gene list—*ERO1B, HNRNPA2B1, HOPX*, and *RHOBTB3*. Combined, these results strongly suggest a type 2 diabetes-relevant role of these genes in beta cells.

## Discussion

In this study, we present an in-depth characterization of the 24 hour transcriptomic response of human pancreatic islets to glucose exposure across cell types and time.

To characterize the effects of glucose exposure on islet cell types through time, we fit a variety of models. Across all cell types, we show that time in culture has a substantial impact on gene expression and, if not experimentally controlled for, will confound differential expression results (e.g., in the case of the BvH model). These findings are an important reminder that islet cell types, especially beta cells, are sensitive to intentional and unintentional experimental perturbations, as we have shown previously (30). An important caveat to remember is that the *in vitro* experimental conditions are not perfect representations of *in vivo* conditions; factors such as time in culture can induce expression changes that must be controlled for. As an example, we found that many of the genes induced by glucose in beta cells also show time in culture effects (Fig. 3), suggesting that primary islets should be analyzed as quickly as possible upon harvesting in order to maintain true islet function.

We find that the number of genes whose expression is associated with glucose exposure plateaus at 8 hours in beta cells, while other cell types showed less consistent trends. Interestingly, for beta cells specifically, we discover that ∼85% of genes differentially expressed at ≥4 hours in LvH models already show expression differences at 1-2 hours, indicating a rapid transcriptomic response to glucose exposure—a finding that would have been missed if we did not serially sample cells over a 24 hour period. In addition, by jointly modeling time and glucose concentration effects on gene expression of cells across all time points, we identify wide-spread multifactorial effects on gene expression in beta cells, whereby genes exhibit expression patterns associated with time in culture, glucose exposure, and time:glucose interaction effects, underscoring the sensitivity of beta cells to glucose exposure and other experimental conditions.

Our work builds on previous glucose stimulation transcriptomic studies in bulk islets (9–11) by characterizing the effects of glucose stimulation on individual islet cell types at multiple time points (Fig. S13-S16, Table S3). From our high-resolution data, we identify specific cell types likely responsible for 489 (10%) of genes previously identified in bulk studies. For 177 genes, we find effects in only one cell type (e.g., *PLIN3* in alpha cells (10), *RASD1* in beta cells (10,11), *DEFB1* in ductal cells (10)), while the remaining 312 genes show effects in multiple cell types (e.g., *GNG4* in alpha and beta cells (10), *EIF5A* in acinar and ductal cells (10)). However, many genes identified as differentially expressed were not shared between our study and previous ones. These differences could be due to a variety of reasons, including (i) differences in islet preparation, (ii) differences in glucose stimulation concentrations, (iii) differences in the duration of glucose stimulation, and (iv) confounding effects of cell type heterogeneity in bulk tissue studies (31).

The availability of cell type expression profiles over 24 hours allowed us to identify modules of genes with similar expression profiles through time in response to glucose stimulation. Interestingly, two beta cell modules were enriched in known type 2 diabetes effector genes. Genes in both modules showed increased expression in high glucose. For module 84, the differences in expression changed over time, particularly after 8 hours. For module 103, the differences in expression remained relatively constant after 4 hours.

Finally, by modeling genetic association summary statistics using genomic features derived from the single-cell data of this study, we nominate additional candidate effector genes for type 2 diabetes, blood glucose, and HbA1c. We identify many genes with a well-defined role in relationship to type 2 diabetes (e.g., *ABCC8, SLC30A8*), including 18 of the 132 type 2 diabetes effector genes from the Type 2 Diabetes Knowledge Portal. Eight of these genes occur in beta cell modules 84 and 103, including four not previously annotated as an effector gene in the type 2 diabetes Knowledge Portal.

In conclusion, one way to help determine the molecular drivers of type 2 diabetes pathogenesis and progression is by understanding the effects of diverse, disease-relevant environmental exposures on gene expression in type 2 diabetes-relevant tissues like pancreatic islets. In this study, we report the effects of sustained glucose exposure on gene expression in islet cell types over the course of 24 hours. Though limited in sample size due to the intensive sampling across time and glucose exposures, these data may provide a relevant window into the consequences of the hyperglycemic conditions that occur as one transitions from impaired glucose tolerance to type 2 diabetes. The results point to coordinated regulation of multiple gene modules in islet cell types in response to glucose stimulation, including several modules that feature known type 2 diabetes genes.

## Materials and Methods

### Ethics statement

The pancreatic islets used in this study were isolated from cadaverous donors whose organs were consented for research. As per the United States’ Office for the Protection of Research Subjects policy, islets obtained from non-living individuals do not fall under the guidelines of human subject research. All experimental protocols performed for this study were approved under National Institutes of Health (NIH) guidelines.

### Islet procurement and processing

We obtained purified human pancreatic islets from two individuals through Prodo Laboratories (Aliso Viejo, CA; Table S1). The purified islets were cultured in PIM(S) complete media (Prodo Laboratories, Aliso Viejo, CA) at a density of 10,000 islet equivalents (IEQs) per 150mm^2^ for 72 hours at 37°C. Islets were packaged and transported to our laboratory at 4°C over a period of 24 hours. Upon receipt, we equilibrated the islets to 37°C for 1 hour in 2.8 mM glucose media prior to downstream processing.

### Single-cell RNA sequencing of glucose-stimulated pancreatic islets

We exposed pancreatic islets to either low (2.8 mM) or high (15 mM) glucose for 24 hours and performed single-cell RNA sequencing (scRNA-seq) at 1, 2, 4, 8, 12, and 24 hour time points. In addition, we performed scRNA-seq on islets at baseline 2.8 mM glucose prior to starting the stimulation experiment (Islet procurement and processing section), which we label as time point 0. To perform the glucose stimulation assay, we plated aliquots of 2,000 IEQs from each donor into four wells of a 12-well plate for each time point (6 plates per donor; two wells for 0 hours since no high glucose exposure) with either low or high glucose in KREBS solution. We incubated the islets at 37°C for the duration of the experiment and sampled wells at their respective time points for scRNA-seq (0, 1, 2, 4, 12, and 24 hours for donor HP18227; 0, 2, 4, 8, 12, and 24 hours for donor HP19208). Each well corresponded to a separate scRNA-seq experiment, resulting in two replicates for each donor, time point, and glucose condition.

For each sampled time point, we dissociated the islet aliquot and performed scRNA-seq. To dissociate the islets, we incubated the 2,000 IEQ aliquot in 1mL Accutase solution (Innovation Cell Technologies, Inc) at 37°C for 10 minutes, washed them with 2 mL PIM(S)TM (Prodo Islet Media, Prodo Laboratory Inc, Irvine CA), incubated the islets for 10 min at 37°C in 2 mL PBS with 4U Dispase I (Roche Diagnostics) / 2U DNase I (ThermoFisher Scientific), washed them once, and resuspended them in PIM(S). We then passed the final cell suspension through a BD 40 mm cell strainer to remove aggregates and assessed the cells for viability and abundance via staining with acridine orange and DAPI (Chemometec Nucleocounter NC-3000). With the filtered suspension, we generated a single-cell mRNA library using either a 10X Genomics SC3’v2 or SC3’v3 chemistry kit according to the manufacturer’s instructions (Table S1). We quantified the barcoded sequencing library with the Quant-IT PicoGreen dsDNA kit (P11496, Invitrogen), diluted cells at 3 nM, and sequenced them on an Illumina HiSeq3000 machine. Across all libraries, we sequenced at an average of 296 million reads per sample.

### Single-cell RNA-seq processing and quality control

We used CellRanger v3.1.0 with default parameters to process and align reads to GRCh38.p13 with Ensembl version 98 transcript definitions (reference file distributed by 10X Genomics), identify cell-containing droplets, and generate cell × gene count matrices.

Next, we used DecontX (32) (implemented in celda v1.14.0 (33)) to (i) remove droplets with a high ambient transcript contamination from the single-cell sequencing experiment and (ii) adjust the raw counts matrix for the ambient expression signature. DecontX requires a gene count matrix of droplets that likely do not contain cells (i.e., “empty droplets”), a count matrix of droplets that likely contain cells, and cell cluster labels. We used the empty droplet count matrix generated by CellRanger, the cell count matrix generated by CellRanger, and cell type labels derived from clustering the cell count matrix with the Seurat v4.3.0 (Hao et al. 2021) single-cell analysis workflow with default parameters, except for using 20 principal components for nearest neighbor calculations. We applied DecontX with default parameters apart from setting the delta parameters to 10 and 30, which represent the prior expectations for the proportion of native and contamination counts, respectively. We removed cell-containing droplets (defined by CellRanger) with >10% ambient contamination as estimated by DecontX from the original CellRanger gene count matrix. Next, we re-ran the DecontX workflow (including Seurat-derived cell type labeling) using the same data and settings apart from removing cells with >10% ambient contamination from the CellRanger cell count matrix. To generate the final cell × gene count matrices, we applied the same ambient contamination filter (removing droplets with >10% ambient contamination) and used the DecontX-adjusted count matrix (Fig. S1A).

We identified and removed multiplets using scrublet v0.2.1 (34), simulating 100,000 multiplets and calculating the multiplet threshold using the threshold_li function from the scikit-image package v0.18.1 (35), initialized using the threshold_otsu function. We removed low-quality cells, defined as cells in which the percentage of counts originating from the mitochondrial genome was >50%. Next, we used an isolation forest (scikit-learn v0.23.2) to remove outlier cells based on (i) the total number of unique molecular identifier (UMI) counts per cell and (ii) the number of genes expressed (≥1 count) per cell. Finally, we verified the reported sex of each sample by generating pseudobulk expression matrices and comparing the expression of *XIST* to the mean expression of all genes on the Y chromosome (both samples were male).

For subsequent analyses, we further processed the data using scanpy v1.6.0 (36). We removed genes that were expressed (≥1 count) in ≤5 cells across the whole dataset (sc.pp.filter_genes with min_cells=5) as well as mitochondrial and ribosomal genes. To account for variable sequencing depth across cells, we normalized UMI counts by the total number of counts per cell, scaled to counts per 10,000 (CP10K; sc.pp.normalise_per_cell), and log-transformed the CP10K expression matrix (ln[CP10K+1]; sc.pp.log1p).

As a final quality control step, we calculated the gene expression correlation between replicates for each donor. Using the cell type annotations (Cell type annotation section), we generated pseudobulk data for each donor replicate, cell type, time point, and glucose condition. We normalized the gene counts to total counts and calculated the Pearson’s correlation coefficient for each donor replicate, cell type, time point, and glucose condition (Fig. S2).

### Cell type annotation

Prior to calculating cell type clusters, we reduced the cell × gene expression matrix to a core set of independent variables using principal component analysis (PCA). First, we selected the 2,000 most variable genes across samples by (i) calculating the most variable genes per sample and (ii) selecting the 2,000 genes that occurred most often across samples (sc.pp.highly_variable_genes with flavor=‘seurat’ and batch_key=sample). After mean centering and scaling the ln(CP10K+1) expression matrix to unit variance, we performed principal component analysis (PCA; sc.tl.pca) using the 2,000 most variable genes. To select the number of PCs for subsequent analyses, we used a scree plot (37) and calculated the “knee/elbow” derived from the variance explained by each PC using the kneedle estimator v0.7.0 (38), selecting 9 PCs. Finally, we used harmony v0.0.5 (39) with default parameters to integrate samples and control for sample specific batch effects prior to clustering.

Using the harmony-adjusted PCs, we calculated clusters using the Leiden graph-based clustering algorithm v0.8.3 (40) (sc.tl.leiden). We generated clusters across a wide range of resolutions to empirically determine the optimal clustering resolution. For each resolution considered, we divided the data into training (2/3 of cells) and test (1/3 of cells) sets. Using the training data, we fit a single layer dense neural network, implemented in keras v2.4.3, to predict cluster identity from the expression of all genes (ln[CP10K+1]). Within the test set, we predicted the cluster label of each cell and calculated the Matthew’s correlation coefficient (MCC) for each cluster, a metric that robustly summarizes all four confusion matrix categories: true positives, false negatives, true negatives, and false positives (41). For the final cluster classifications, we chose the largest resolution where the minimum MCC across all clusters was >0.75, selecting a resolution of 0.25 (8 clusters, Fig. S1B-C).

To determine the cell type identity of clusters, we used well-established marker genes for the cell types common to islets (Fig. S1D): *GCG* (alpha cells), *INS* (beta cells), *PPY* (gamma cells), *SST* (delta cells), *PRSS1* (acinar cells), *KRT19* (ductal cells), *COL1A1* (stellate cells), and *CD68* (macrophages).

### Time interpolation

We derived interpolated time from sampled time points (t=1h, 2h, 4h, …, 24h) to better model the cellular state of each cell at each time point, assuming that cells captured at each time point are not at a uniform cellular state in their response to time in culture and/or glucose exposure but rather are distributed across cellular response phases. Prior to calculating interpolated time, we removed the 0 hour time point as there was no high glucose treatment at this time point. For each cell type and glucose condition, we calculated the *n* nearest neighbors based on gene expression values (ln[CP10K+1]) using scvelo v0.2.4 (scv.pp.neighbors) and derived interpolated time, 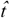, for each cell using the following formula:

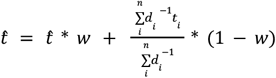

where *t* is the target cell’s sampled time point, *w* is a weight coefficient for *t, d*_*i*_ is the distance of the target cell to neighboring cell *i*, and *t*_*i*_ is the sampled time point of neighboring cell *i*. To select values for *n* and *w*, we performed an exhaustive grid search to evaluate the effect of all combinations of *n* and *w* on the stability of 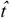, where *n* ranged from 0 to 150 in intervals of 5 and *w* ranged from 0 to 1 in intervals of 0.25. For each value of *w*, we considered all values of *n* and compared 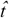 to 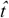 derived from the previous *n* value using mean squared error (MSE) as a stability metric (Fig. S8A). Across all values of *w*, we found 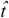 was stable at *n*≥75 and selected a value of *n*=75 for further analyses. To determine the value of *w*, we set *n*=75, calculated 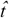 for *w* values running from 0 to 1 in intervals of 0.25, standardized 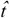, and evaluated the effect of different 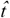 derivations on the differential gene expression results for the time:glucose interaction model (Differential gene expression analysis section), comparing the signed -log_10_(*P*-values) for each 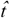 derivation to the results of using *t* (the sampled time point; Fig. S8B). We found the results were highly correlated across all values of *w*. For subsequent analysis, we selected a value of 0.25 for *w* since the results were extremely concordant with *t*, yet showed a slight increase in power. We calculated the final interpolated time values using *n*=75 and *w*=0.25 and standardized interpolated time values within each glucose condition and cell type to a 0-1 scale.

### Differential gene expression analysis

For each cell type, we performed differential gene expression (DGE) analysis (both discrete time point DGE models and continuous time DGE models) using MAST v1.20.0, a two-part, generalized linear model with a logistic regression component for the discrete process (i.e., a gene is expressed or not) and linear regression component for the continuous process (i.e., the expression level) (42). Briefly, for gene *i*, and cell *k*, let *Z*_*ki*_ indicate whether gene *i* is expressed in cell *k* and *Y*_*ki*_ denote the ln(CP10K+1) normalized gene expression. We tested for association using the two-part regression model:

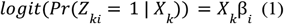

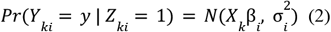

where *X*_*k*_ are the predictor variables for cell *k* and *β*_*i*_ is the vector of fixed effect regression coefficients.

Across all DGE models, we included as fixed effect covariates cell complexity (i.e., the number of genes detected per cell) to control for unobserved nuisance variation (e.g., cell size) (42) and participant identifiers to control for pseudoreplication bias (43). We also included additional fixed effect variables depending on the specific model (see subsequent paragraphs).

For the discrete time DGE models, we fit separate models for each cell type and time point. We considered three models. First, in the “basal-versus-high” (BvH) model, we compared basal cells (0 hours, 2.8 mM glucose) to cells exposed to high glucose (15 mM) across time (e.g., basal versus 1 hour of high glucose exposure, basal versus 2 hours of high glucose exposure) by including a fixed effect “glucose status” variable (0 = basal/low, 1 = high). Second, in the “basal-versus-low” (BvL) model, we compared basal cells (0 hours, 2.8 mM glucose) to cells maintained in low glucose (2.8 mM) across time (e.g., basal versus 1 hour of low glucose exposure, basal versus 2 hours of low glucose exposure) by including a fixed effect “time point” variable (0 = basal at 0 hours, 1 = other time points). Third, in the “low-versus-high glucose” (LvH) model, we compared cells exposed to low glucose (2.8 mM) to cells exposed to high glucose (15 mM) at each time point (e.g., 1 hour of low glucose exposure versus 1 hour of high glucose exposure, 2 hours of low glucose exposure versus 2 hours of high glucose exposure) by including a fixed effect “glucose status” variable (0 = low, 1 = high).

For the continuous time DGE models, we fit separate models for each cell type, jointly analyzing cells across all glucose exposures and time points, excluding the 0 hour time point. We considered three different models: (i) a “continuous time” model to test for time effects, (ii) a “glucose” model to test for glucose effects, and (iii) a “time:glucose interaction” model to test for an interaction effect between time and glucose concentration. For all three models, we included glucose concentration (0 = low, 1 = high) and continuous time as fixed effect variables (in addition to cell complexity and participant identifiers). For the “time:glucose interaction” model, we used the same model but added an additional time:glucose concentration interaction term.

Finally, for each model, we controlled for the number of tests performed with each cell type using the Benjamini-Hochberg procedure (44) and *P*-values obtained from the hurdle model, derived from the summed *χ*^*2*^ null distributions of the discrete (*Z*_*i*_) and continuous (*Y*_*i*_) components, as described in Finak et al. (42). To increase the speed of each test, prior to fitting models for each cell type, we removed genes with a median CP10K<1.

### Identification of interpolated time and glucose gene expression modules

For each cell type, we clustered the expression profiles (ln[CP10K+1]) of all genes associated (FDR<5%) with any of the DGE models apart from the BvH model (Differential gene expression analysis section). Before generating clusters, we split the data based on the glucose exposure condition (low or high) and removed the 0 hour time point as there was no high glucose treatment at this time point. For each gene, we fit a local polynomial regression using skmisc v0.1.4 to model gene expression as a function of interpolated time, resulting in two regressions—one for low glucose exposure and one for high glucose exposure. From each regression, we generated a predicted gene expression × interpolated time matrix using interpolated time values ranging from 0 to 1 in steps of 0.05 in order to gain a more fine-grain characterization of expression over time than would be possible using sampled time.

We clustered the gene expression × interpolated time matrices (one for each glucose condition) using the Dirichlet process Gaussian process (DPGP) package v0.1 (45). In order to ensure that the final clusters assignments were stable, we used a consensus clustering approach (46) where we (i) ran DPGP over a variety of parameter configurations, (ii) calculated a consensus matrix, defined as the frequency at which two genes cluster with each other across all parameter configurations, and (iii) clustered the consensus matrix using *k*-means (kmeans function in R). For the DPGP parameter configurations, we considered 45 different parameter configurations, ranging (i) the alpha parameter, which tunes the number of clusters, from 0.001 to 10.0 and (ii) the num_empty_clusters parameter, that governs the number of empty clusters introduced at each iteration of the clustering algorithm, from 2 to 16. For each parameter configuration, we merged the DPGP clusters across low and high glucose by (i) averaging the similarity matrices, (ii) concatenating the cluster assignments and the corresponding log-likelihood values, and (iii) re-running DPGP using the merged information from steps i-ii. For the *k*-means clustering of the consensus matrix, we started at *k*=2 and iteratively increased *k* by 1 until ≥1 cluster had a minimum co-clustering frequency ≥0.75. We removed all genes in the cluster(s) with a minimum co-clustering frequency ≥0.75 from the consensus matrix, subtracted the number of removed clusters from *k*, and repeated this process—iteratively increasing *k* and removing clusters with a co-clustering frequency ≥0.75 until no genes remained in the consensus matrix. For final module assignments, we removed clusters with <5 genes.

To determine T2D-relevant expression modules, we tested the modules for enrichment (hypergeometric test; phyper function in R) of T2D effector genes in the genes constituting the module. For T2D effector gene definitions, we used the Mahajan and McCarthy effector gene predictions from the T2D knowledge portal (https://t2d.hugeamp.org/method.html?trait=t2d&dataset=mccarthy). We controlled for the number of tests performed for each cell type using the Benjamini-Hochberg procedure.

### Gene ontology enrichment and clustering analysis

For the gene expression modules that were enriched in T2D-relevant genes (Identification of interpolated time and glucose gene expression modules section), we performed gene ontology (GO) term enrichment calculations across all ontology classes (biological function, cellular component, molecular function) using the enrichGO method from clusterProfiler v4.0.5 (47). We removed GO terms with only one gene overlap and controlled for the number of tests within each module using the Benjamini-Hochberg procedure. To aid in visualization, we simplified the GO term results by clustering the enriched GO terms (FDR<5%) for each ontology across all modules using the simplifyGO function from simplifyEnrichment v4.0.5 (47). We used (i) the relevance semantic similarity measurement (48), implemented in GOSemSim v2.18.1 (49), as a distance metric for clustering and (ii) affinity propagation, implemented in apcluster v1.4.10 (50), as the clustering algorithm. We report the full set of enriched terms in Fig. S11; for the main text, we report terms with ≥2 gene overlap with the expression modules (cellular component terms are missing from these plots as we identified no enriched terms at an FDR<5%).

### Nomination of candidate effector genes for T2D and T2D-related traits

We nominated candidate effector genes using the Polygenic Priority Score (PoPS) method v0.2 (29). Briefly, the PoPS method requires summary statistics from a genetic association study and a genomic feature matrix across genes (gene × feature matrix). For the summary statistics, we used publicly available summary statistics for T2D (7), blood glucose (8), and glycated hemoglobin (8). We constructed a genomic feature matrix from the single-cell data presented in this study as performed in Weeks et al. (29), including (i) principal component gene weights across all cells and within cells for each cell type, (ii) test statistics for cell type marker genes (Welch’s *t*-test; including the test statistic and a binary indicator for up- and down-regulated genes with FDR<5%), and (iii) average expression within each cell type. In addition, we also included (i) cell type gene expression specificity values calculated using CELLEX v1.2.2 (51) and (ii) test statistics from all models but the BvH model (Differential gene expression analysis section, including the test statistic and a binary indicator for up- and down-regulated genes with FDR<5%). To calculate *P*-values, we permuted the feature matrix gene identifiers 10,000 times and re-ran PoPS. We pooled the permuted PoP scores across all genes and calculated empirical *P-*values, defined as (*r*+1) / (*n*+1), where *r* is the number of permuted PoP scores greater than the observed and *n* is the total number of permutations (52). Finally, we controlled the false discovery rate across all genes considered using the Benjamini-Hochberg procedure.

### Comparison with results of previous bulk islet transcriptome studies

Previous bulk islet transcriptomic studies report genes associated with glucose stimulation using a study design similar to the LvH model of this study (9–11). We compared the results of all glucose-relevant models (discrete models: LvH all time points; continuous models: glucose, time:glucose interaction) to studies where complete lists of differentially expressed genes were available: Ottosson-Laakso et al. (10) (5.5 mM vs. 18.9 mM glucose for 24 hours in normoglycemic and hyperglycemic donors) and Hall et al. (11) (5.6 mM vs. 19 mM for 48 hours). The results from these comparisons are recorded in Fig. S13-S16 and Table S3.

## Acknowledgments

Author contributions: C.G., L.L.B., M.R.E., F.S.C., and D.L.T. designed research; C.G., L.L.B., A.J.S., C.C.R., M.R.E., F.S.C., and D.L.T. performed research; C.G., H.J.T., T.Y., N.N., and D.L.T analyzed data; F.S.C. and D.L.T. supervised the study; and C.G., L.L.B., H.J.T., F.S.C., and D.L.T. wrote the paper.

This research was supported in part by the United States’ National Institutes of Health grant ZIA-HG000024 (to F.S.C.), the Gates Cambridge Trust (to H.J.T.), and the NIH Oxford-Cambridge scholars’ program (to H.J.T.). We thank Chad Krilow for supporting early parts of the work presented in this study.

## References

1. International Diabetes Federation. IDF Diabetes Atlas. 10th ed. Brussels, Belgium: International Diabetes Federation; 2021 Jan.

2. Krentz NAJ, Gloyn AL. Insights into pancreatic islet cell dysfunction from type 2 diabetes mellitus genetics. Nat Rev Endocrinol. 2020 Apr;16(4):202–12.

3. Ahmad E, Lim S, Lamptey R, Webb DR, Davies MJ. Type 2 diabetes. Lancet. 2022 Nov 19;400(10365):1803–20.

4. Ashcroft FM, Rorsman P. Diabetes mellitus and the β cell: the last ten years. Cell. 2012 Mar 16;148(6):1160–71.

5. Gromada J, Chabosseau P, Rutter GA. The α-cell in diabetes mellitus. Nat Rev Endocrinol. 2018 Dec;14(12):694–704.

6. Gao R, Yang T, Zhang Q. δ-Cells: The Neighborhood Watch in the Islet Community. Biology (Basel). 2021 Jan 21;21(10).

7. Mahajan A, Taliun D, Thurner M, Robertson NR, Torres JM, Rayner NW, et al. Fine-mapping type 2 diabetes loci to single-variant resolution using high-density imputation and islet-specific epigenome maps. Nat Genet. 2018 Nov;50(11):1505–13.

8. Sinnott-Armstrong N, Tanigawa Y, Amar D, Mars N, Benner C, Aguirre M, et al. Genetics of 35 blood and urine biomarkers in the UK Biobank. Nat Genet. 2021 Feb;53(2):185–94.

9. Taneera J, Fadista J, Ahlqvist E, Atac D, Ottosson-Laakso E, Wollheim CB, et al. Identification of novel genes for glucose metabolism based upon expression pattern in human islets and effect on insulin secretion and glycemia. Hum Mol Genet. 2015 Apr 1;24(7):1945–55.

10. Ottosson-Laakso E, Krus U, Storm P, Prasad RB, Oskolkov N, Ahlqvist E, et al. Glucose-Induced Changes in Gene Expression in Human Pancreatic Islets: Causes or Consequences of Chronic Hyperglycemia. Diabetes. 2017 Dec;66(12):3013–28.

11. Hall E, Dekker Nitert M, Volkov P, Malmgren S, Mulder H, Bacos K, et al. The effects of high glucose exposure on global gene expression and DNA methylation in human pancreatic islets. Mol Cell Endocrinol. 2018 Sep 5;472:57–67.

12. Kaiser N, Leibowitz G, Nesher R. Glucotoxicity and beta-cell failure in type 2 diabetes mellitus. J Pediatr Endocrinol Metab. 2003 Jan;16(1):5–22.

13. Guardado Mendoza R, Perego C, Finzi G, La Rosa S, Capella C, Jimenez-Ceja LM, et al. Delta cell death in the islet of Langerhans and the progression from normal glucose tolerance to type 2 diabetes in non-human primates (baboon, Papio hamadryas). Diabetologia. 2015 Aug;58(8):1814–26.

14. Proks P, Arnold AL, Bruining J, Girard C, Flanagan SE, Larkin B, et al. A heterozygous activating mutation in the sulphonylurea receptor SUR1 (ABCC8) causes neonatal diabetes. Hum Mol Genet. 2006 Jun 1;15(11):1793–800.

15. Babenko AP, Polak M, Cavé H, Busiah K, Czernichow P, Scharfmann R, et al. Activating mutations in the ABCC8 gene in neonatal diabetes mellitus. N Engl J Med. 2006 Aug 3;355(5):456–66.

16. Scott LJ, Mohlke KL, Bonnycastle LL, Willer CJ, Li Y, Duren WL, et al. A genome-wide association study of type 2 diabetes in Finns detects multiple susceptibility variants. Science. 2007 Jun 1;316(5829):1341–5.

17. Flannick J, Thorleifsson G, Beer NL, Jacobs SBR, Grarup N, Burtt NP, et al. Loss-of-function mutations in SLC30A8 protect against type 2 diabetes. Nat Genet. 2014 Apr;46(4):357–63.

18. Stijnen P, Tuand K, Varga TV, Franks PW, Aertgeerts B, Creemers JWM. The association of common variants in PCSK1 with obesity: a HuGE review and meta-analysis. Am J Epidemiol. 2014 Dec 1;180(11):1051–65.

19. Philippe J, Stijnen P, Meyre D, De Graeve F, Thuillier D, Delplanque J, et al. A nonsense loss-of-function mutation in PCSK1 contributes to dominantly inherited human obesity. Int J Obes (Lond). 2015 Feb;39(2):295–302.

20. Stijnen P, Ramos-Molina B, O’Rahilly S, Creemers JWM. PCSK1 mutations and human endocrinopathies: from obesity to gastrointestinal disorders. Endocr Rev. 2016 May 17;37(4):347–71.

21. Nead KT, Li A, Wehner MR, Neupane B, Gustafsson S, Butterworth A, et al. Contribution of common non-synonymous variants in PCSK1 to body mass index variation and risk of obesity: a systematic review and meta-analysis with evidence from up to 331 175 individuals. Hum Mol Genet. 2015 Jun 15;24(12):3582–94.

22. Swisa A, Avrahami D, Eden N, Zhang J, Feleke E, Dahan T, et al. PAX6 maintains β cell identity by repressing genes of alternative islet cell types. The Journal of Clinical Investigation. 2017 Jan 3;

23. Gosmain Y, Katz LS, Masson MH, Cheyssac C, Poisson C, Philippe J. Pax6 is crucial for β-cell function, insulin biosynthesis, and glucose-induced insulin secretion. Mol Endocrinol. 2012 Apr;26(4):696–709.

24. Yasuda T, Kajimoto Y, Fujitani Y, Watada H, Yamamoto S, Watarai T, et al. PAX6 mutation as a genetic factor common to aniridia and glucose intolerance. Diabetes. 2002 Jan;51(1):224–30.

25. Boortz KA, Syring KE, Dai C, Pound LD, Oeser JK, Jacobson DA, et al. G6PC2 modulates fasting blood glucose in male mice in response to stress. Endocrinology. 2016 Aug;157(8):3002–8.

26. Pound LD, Oeser JK, O’Brien TP, Wang Y, Faulman CJ, Dadi PK, et al. G6PC2: a negative regulator of basal glucose-stimulated insulin secretion. Diabetes. 2013 May;62(5):1547–56.

27. Reiling E, van ‘t Riet E, Groenewoud MJ, Welschen LMC, van Hove EC, Nijpels G, et al. Combined effects of single-nucleotide polymorphisms in GCK, GCKR, G6PC2 and MTNR1B on fasting plasma glucose and type 2 diabetes risk. Diabetologia. 2009 Sep;52(9):1866–70.

28. Bouatia-Naji N, Rocheleau G, Van Lommel L, Lemaire K, Schuit F, Cavalcanti-Proença C, et al. A polymorphism within the G6PC2 gene is associated with fasting plasma glucose levels. Science. 2008 May 23;320(5879):1085–8.

29. Weeks EM, Ulirsch JC, Cheng NY, Trippe BL, Fine RS, Miao J, et al. Leveraging polygenic enrichments of gene features to predict genes underlying complex traits and diseases. medRxiv. 2020 Sep 10;

30. Bonnycastle LL, Gildea DE, Yan T, Narisu N, Swift AJ, Wolfsberg TG, et al. Single-cell transcriptomics from human pancreatic islets: sample preparation matters. Biol Methods Protoc. 2020 Jan 16;5(1):bpz019.

31. Taylor DL, Jackson AU, Narisu N, Hemani G, Erdos MR, Chines PS, et al. Integrative analysis of gene expression, DNA methylation, physiological traits, and genetic variation in human skeletal muscle. Proc Natl Acad Sci USA. 2019 May 28;116(22):10883–8.

32. Yang S, Corbett SE, Koga Y, Wang Z, Johnson WE, Yajima M, et al. Decontamination of ambient RNA in single-cell RNA-seq with DecontX. Genome Biol. 2020 Mar 5;21(1):57.

33. Wang Z, Yang S, Koga Y, Corbett SE, Shea CV, Johnson WE, et al. Celda: a Bayesian model to perform co-clustering of genes into modules and cells into subpopulations using single-cell RNA-seq data. NAR Genom Bioinform. 2022 Sep 13;4(3):qac066.

34. Wolock SL, Lopez R, Klein AM. Scrublet: Computational Identification of Cell Doublets in Single-Cell Transcriptomic Data. Cell Syst. 2019 Apr 24;8(4):281–291.e9.

35. van der Walt S, Schönberger JL, Nunez-Iglesias J, Boulogne F, Warner JD, Yager N, et al. scikit-image: image processing in Python. PeerJ. 2014 Jun 19;2:e453.

36. Wolf FA, Angerer P, Theis FJ. SCANPY: large-scale single-cell gene expression data analysis. Genome Biol. 2018 Feb 6;19(1):15.

37. Cattell RB. The scree test for the number of factors. Multivariate Behav Res. 1966 Apr 1;1(2):245–76.

38. Satopaa V, Albrecht J, Irwin D, Raghavan B. Finding a “kneedle” in a haystack: detecting knee points in system behavior. 2011 31st International Conference on Distributed Computing Systems Workshops. IEEE; 2011. p. 166–71.

39. Polański K, Young MD, Miao Z, Meyer KB, Teichmann SA, Park J-E. BBKNN: fast batch alignment of single cell transcriptomes. Bioinformatics. 2020 Feb 1;36(3):964–5.

40. Traag VA, Waltman L, van Eck NJ. From Louvain to Leiden: guaranteeing well-connected communities. Sci Rep. 2019 Mar 26;9(1):5233.

41. Chicco D, Jurman G. The advantages of the Matthews correlation coefficient (MCC) over F1 score and accuracy in binary classification evaluation. BMC Genomics. 2020 Jan 2;21(1):6.

42. Finak G, McDavid A, Yajima M, Deng J, Gersuk V, Shalek AK, et al. MAST: a flexible statistical framework for assessing transcriptional changes and characterizing heterogeneity in single-cell RNA sequencing data. Genome Biol. 2015 Dec 10;16:278.

43. Zimmerman KD, Espeland MA, Langefeld CD. A practical solution to pseudoreplication bias in single-cell studies. Nat Commun. 2021 Feb 2;12(1):738.

44. Benjamini Y, Hochberg Y. Controlling the false discovery rate: A practical and powerful approach to multiple testing. Journal of the Royal Statistical Society Series B (Methodological). 1995;57(1):289–300.

45. McDowell IC, Manandhar D, Vockley CM, Schmid AK, Reddy TE, Engelhardt BE. Clustering gene expression time series data using an infinite Gaussian process mixture model. PLoS Comput Biol. 2018 Jan 16;14(1):e1005896.

46. Steinley D. Stability analysis in K-means clustering. Br J Math Stat Psychol. 2008 Nov;61(Pt 2):255–73.

47. Wu T, Hu E, Xu S, Chen M, Guo P, Dai Z, et al. clusterProfiler 4.0: A universal enrichment tool for interpreting omics data. Innovation (Camb). 2021 Aug 28;2(3):100141.

48. Schlicker A, Domingues FS, Rahnenführer J, Lengauer T. A new measure for functional similarity of gene products based on Gene Ontology. BMC Bioinformatics. 2006 Jun 15;7:302.

49. Yu G, Li F, Qin Y, Bo X, Wu Y, Wang S. GOSemSim: an R package for measuring semantic similarity among GO terms and gene products. Bioinformatics. 2010 Apr 1;26(7):976–8.

50. Frey BJ, Dueck D. Clustering by passing messages between data points. Science. 2007 Feb 16;315(5814):972–6.

51. Timshel PN, Thompson JJ, Pers TH. Genetic mapping of etiologic brain cell types for obesity. eLife. 2020 Sep 21;9.

52. North BV, Curtis D, Sham PC. A note on the calculation of empirical P values from Monte Carlo procedures. Am J Hum Genet. 2002 Aug;71(2):439–41.

