## Supplemental Figures for "Single-cell transcriptomic profiling of human pancreatic islets reveals genes responsive to glucose exposure over 24 hours"

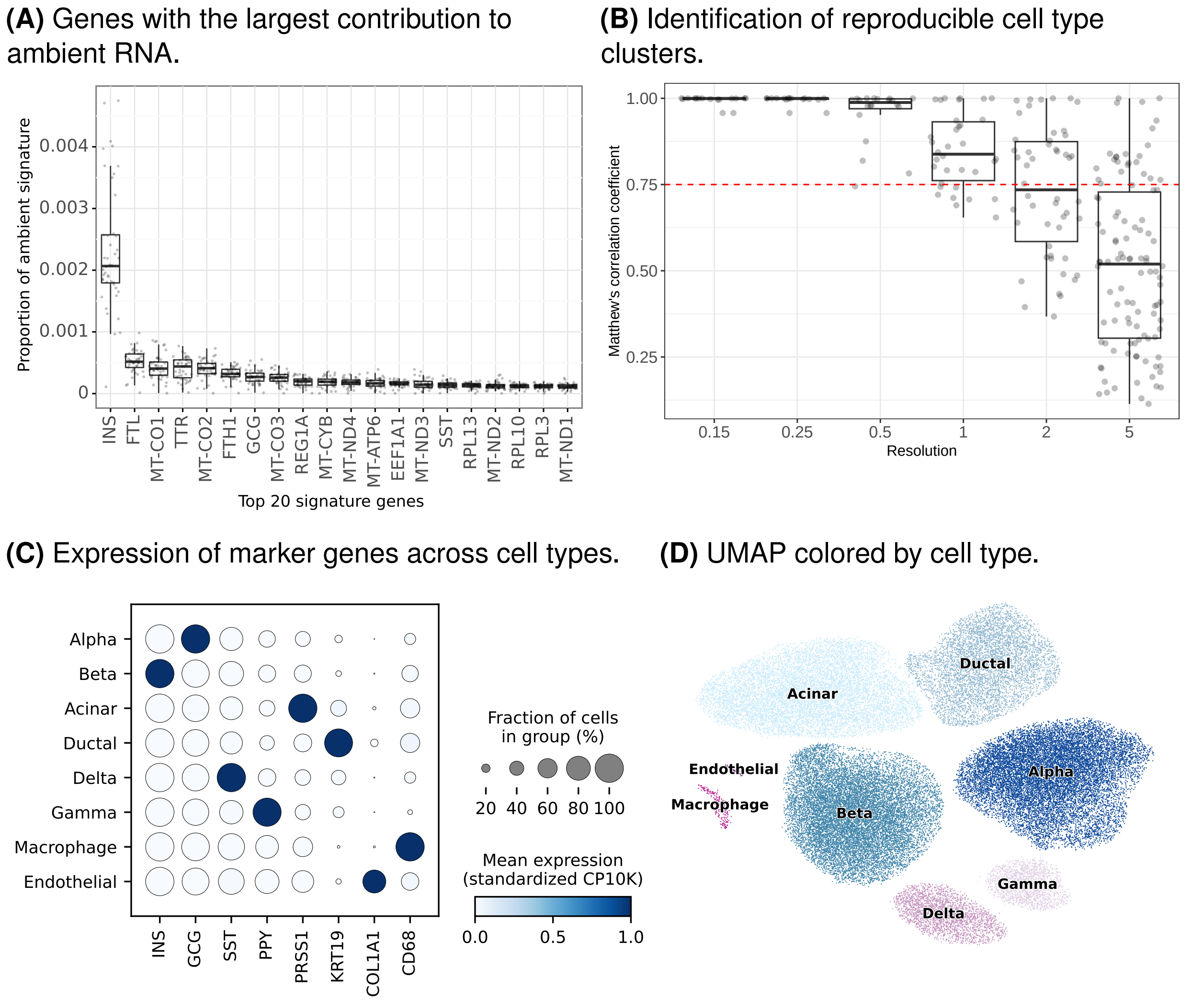


**Fig. S1. Summary of single-cell RNA-seq data and cell type clusters.** (A) 20 genes (x-axis) with the largest contribution to the ambient RNA signature (y-axis) across samples. (B) Distribution of cluster (points) predictability measured using Matthew's correlation coefficient (y-axis) across different clustering resolution values (x-axis). Dashed line at 0.75. (C) Expression of cell type marker genes (x-axis) across final clusters (y-axis). (D) Uniform manifold approximation and projection (UMAP) dimensions colored by cell type.

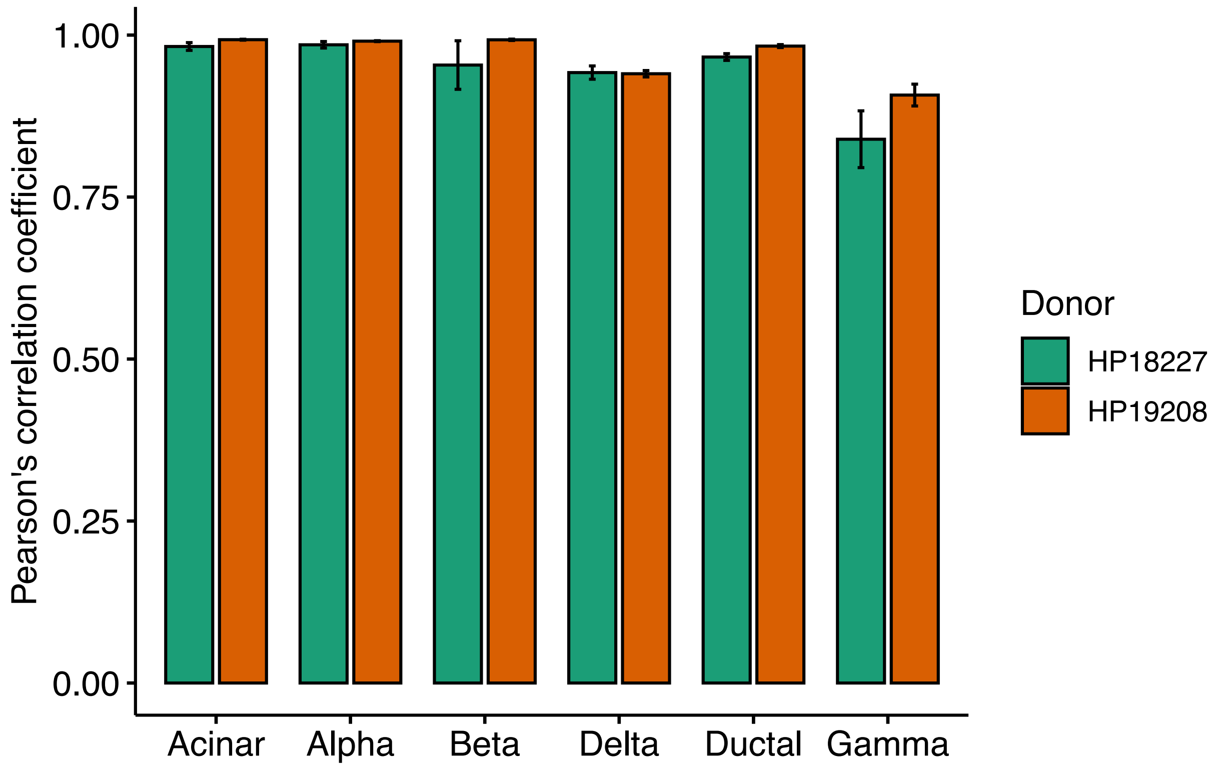


**Fig. S2. Correlation between replicates**. Average correlation (y-axis) of gene expression between replicates of each donor (color) for each cell type (x-axis) at each time point and glucose condition. Bars depict standard error.

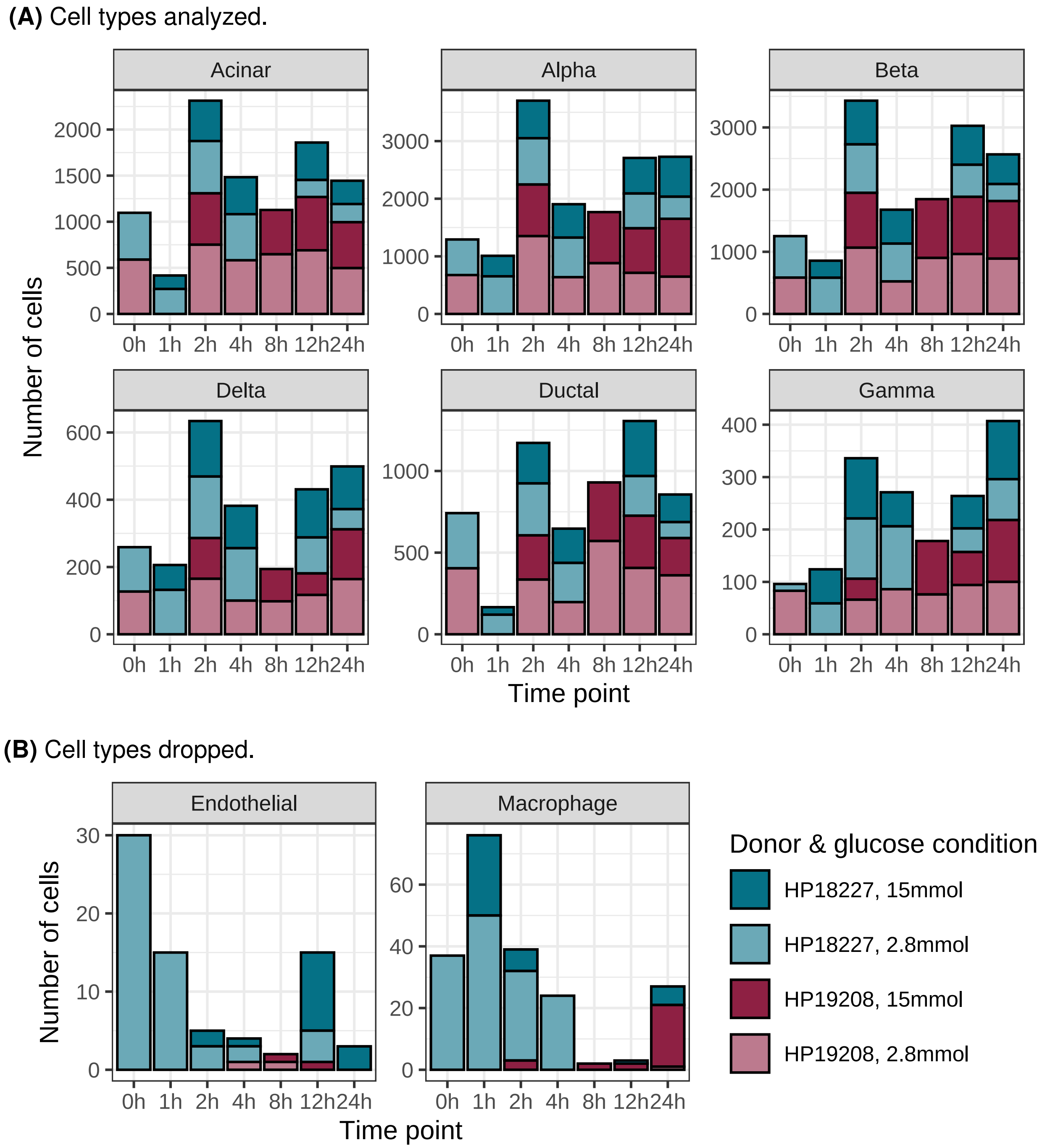


**Fig. S3. Donor representation within cell types**. Number of cells (y-axis) from each donor (colors) across cell types (facets) at each time point (x-axis). Darker shades represent donor cells from the higher glucose exposure, and lighter shades represent lower glucose exposure. (A) Cell types considered in analyses. (B) Low-frequency cell types dropped from analyses.

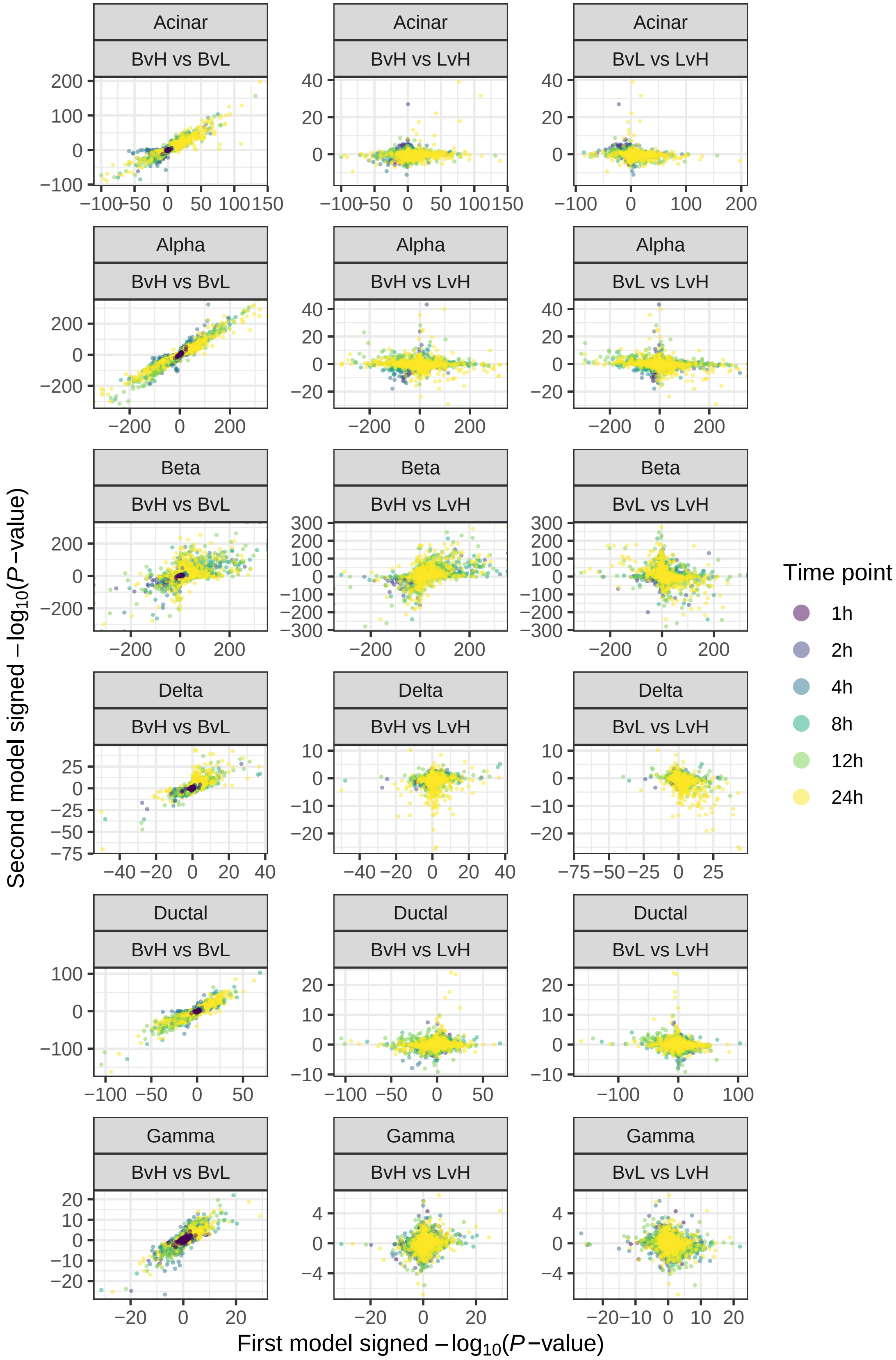


**Fig. S4. Comparison of differential gene expression results across discrete time models.** Signed -$\log_{10}$($P$-values) of BvL, BvH, and LvH models for each cell type. X-axis corresponds to the first model listed in facets, y-axis corresponds to the second model listed in facets.

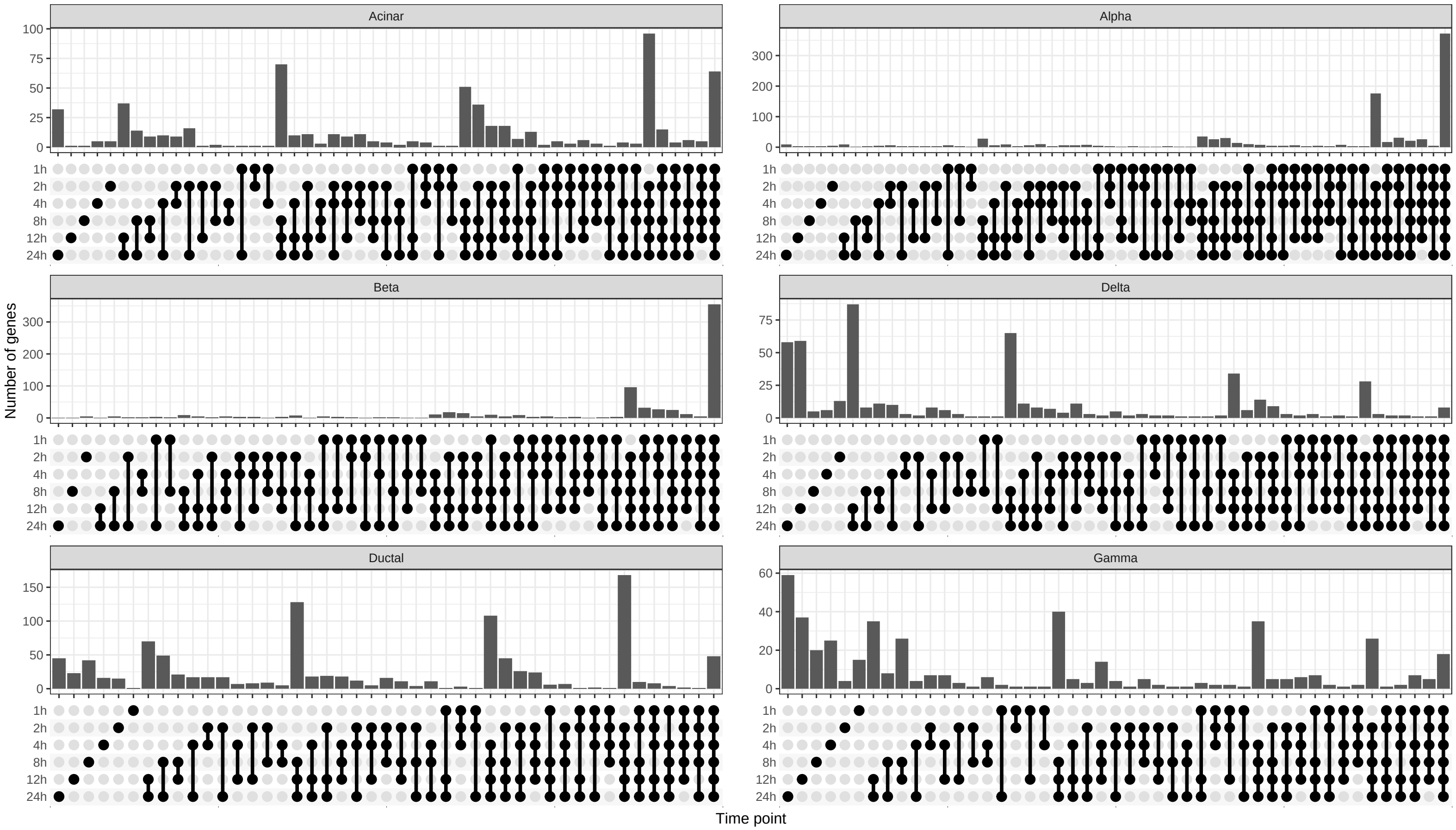


**Fig. S5. Overlap of differentially expressed genes in BvL models across time points.** Number of associated genes (FDR<5%; y-axis) shared between time points (x-axis) in the BvL model within each cell type.

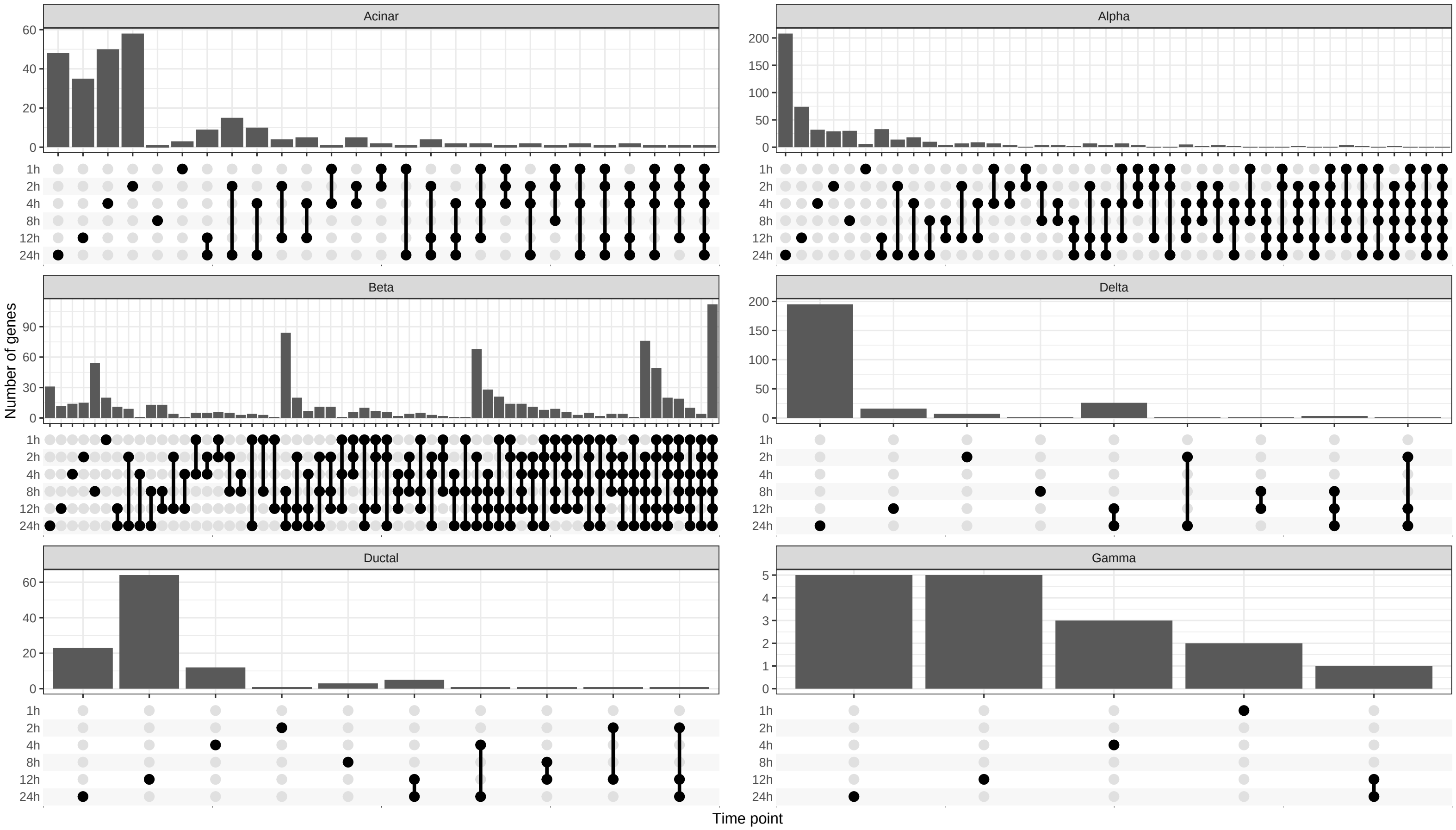


**Fig. S6. Overlap of differentially expressed genes in LvH models across time points.** Number of associated genes (FDR<5%; y-axis) shared between time points (x-axis) in the LvH model within each cell type.

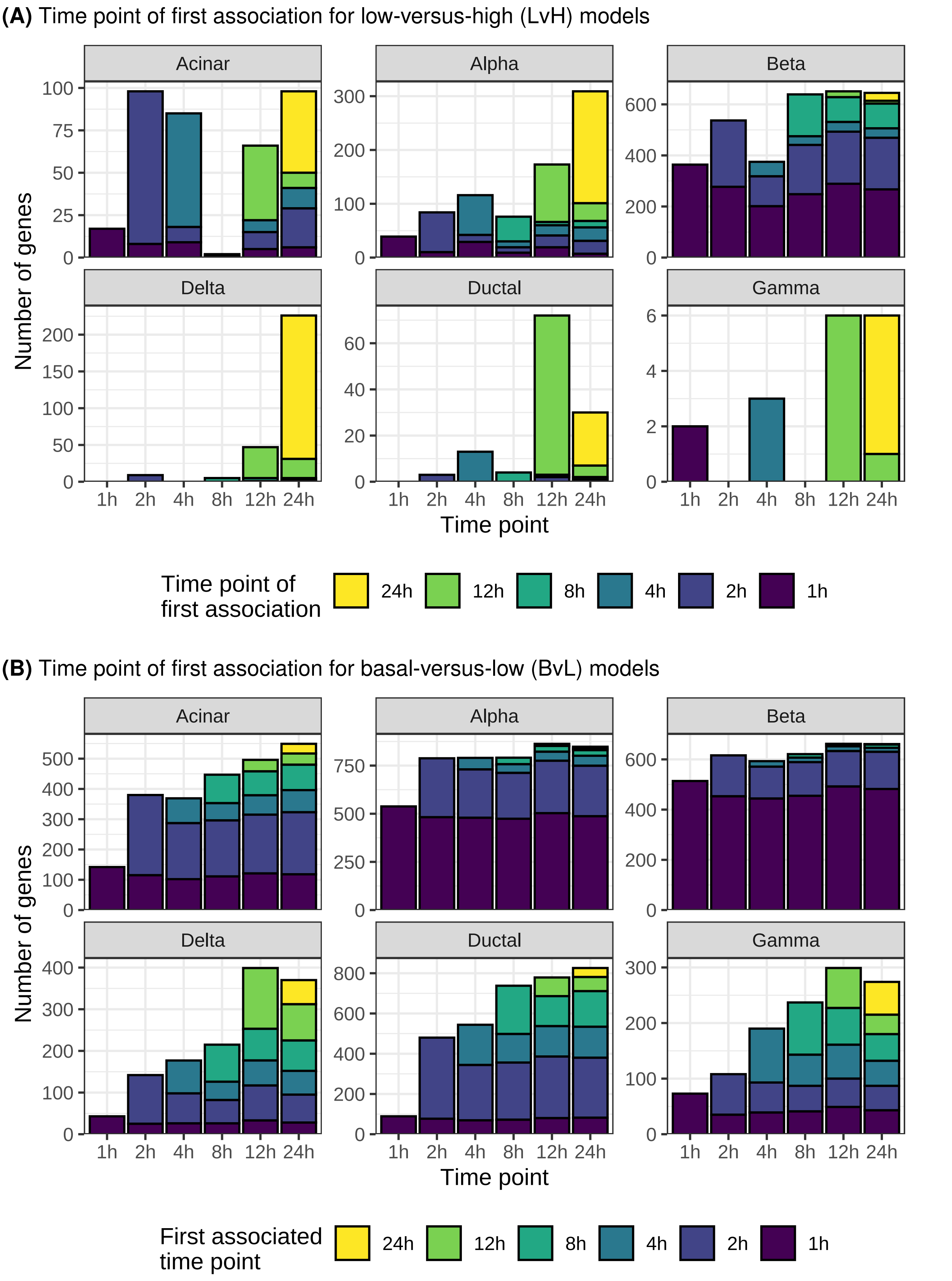


**Fig. S7. Time point of first association for LvH and BvL models.** Number of associated genes (FDR<5%; y-axis) for each time point (x-axis) across cell types. Color denotes the time point where the gene was first identified as differentially expressed. (A) LvH model. (B) BvL model.

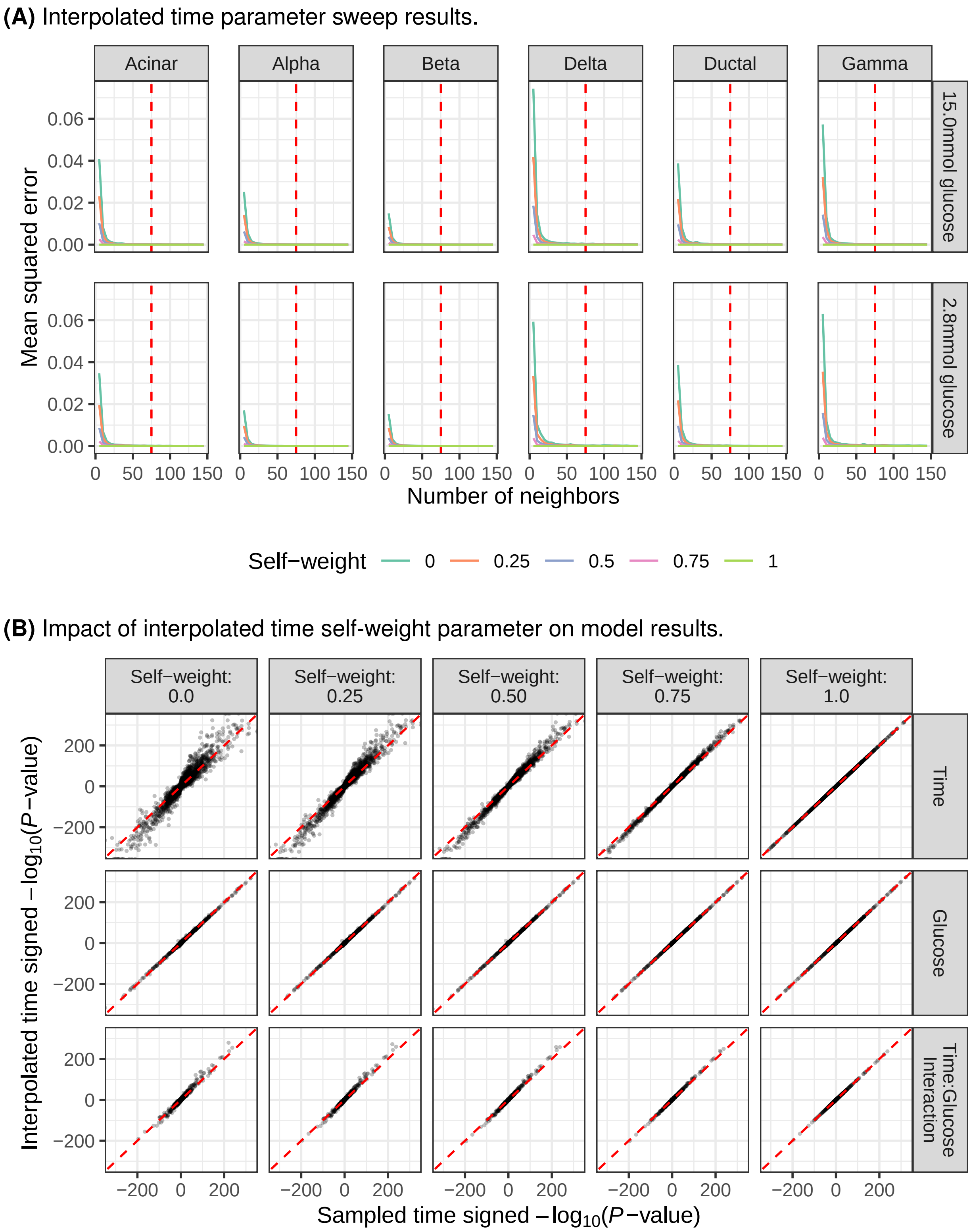


**Fig. S8. Evaluation of parameters used to calculate interpolated time.** (A) Mean squared error (y-axis) calculated by comparing the per-cell interpolated time for each "number of neighbors" value (x-axis) to the per-cell interpolated time of the previous number of neighbors. Line colors correspond to different self-weight values. Red dashed line at 75. (B) Comparison of signed -$\log_{10}$($P$-values) across continuous models (row facets) when using sampled time (x-axis) and interpolated time (y-axis) across self-weight values (column facets). Number of neighbors set to 75 for interpolated time calculation.

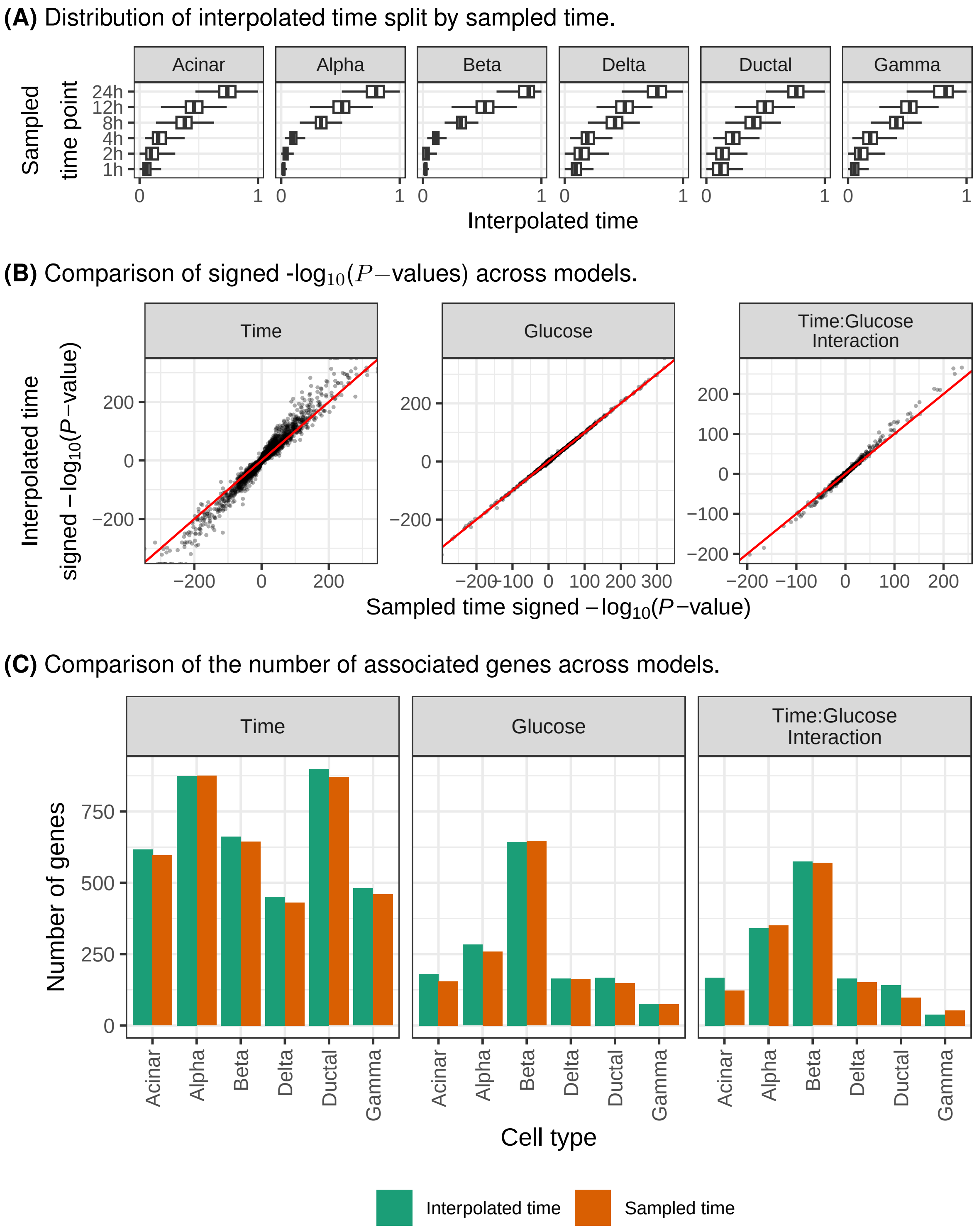


**Fig. S9. Comparison of differential gene expression results for continuous models using interpolated or sampled time.** (A) Distribution of interpolated time (x-axis) for cells from each sampled time point (y-axis) across cell types. (B) Comparison of signed -$\log_{10}$($P$-values) of differential gene expression results for continuous models using sampled time (x-axis) and interpolated time (y-axis). (C) Number of associated genes (FDR$<$5%; y-axis) for each cell type (x-axis) across continuous models (facets). Color indicates if the model used interpolated or sampled time.

**(A)** Intersection of continuous time models with discrete models

**(B)** Intersection of continuous glucose models with discrete models

**(C)** Intersection of continuous time:glucose interaction models with discrete models

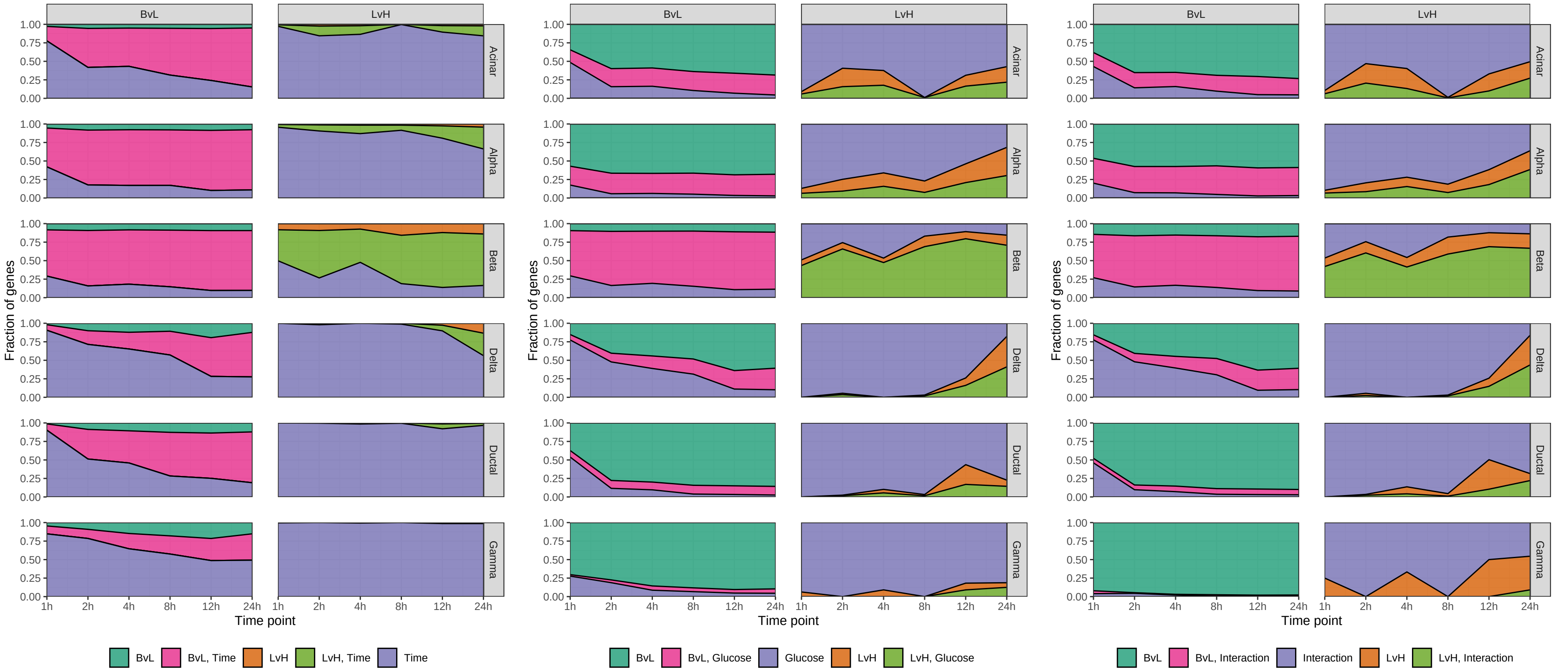


**Fig. S10. Intersection of differential gene expression results from continuous models with discrete models.** Fraction of associated genes (FDR<5%; y-axis) shared between continuous models and discrete models (column facets) across cell types (row facets) and discrete model time points (x-axis). Color denotes the combination of models that genes belong to. (A) Continuous time model (abbreviated as “Time” in color legend). (B) Continuous glucose models (abbreviated as “Glucose” in color legend). (C) Continuous time:glucose interaction model (abbreviated as “Interaction” in color legend).

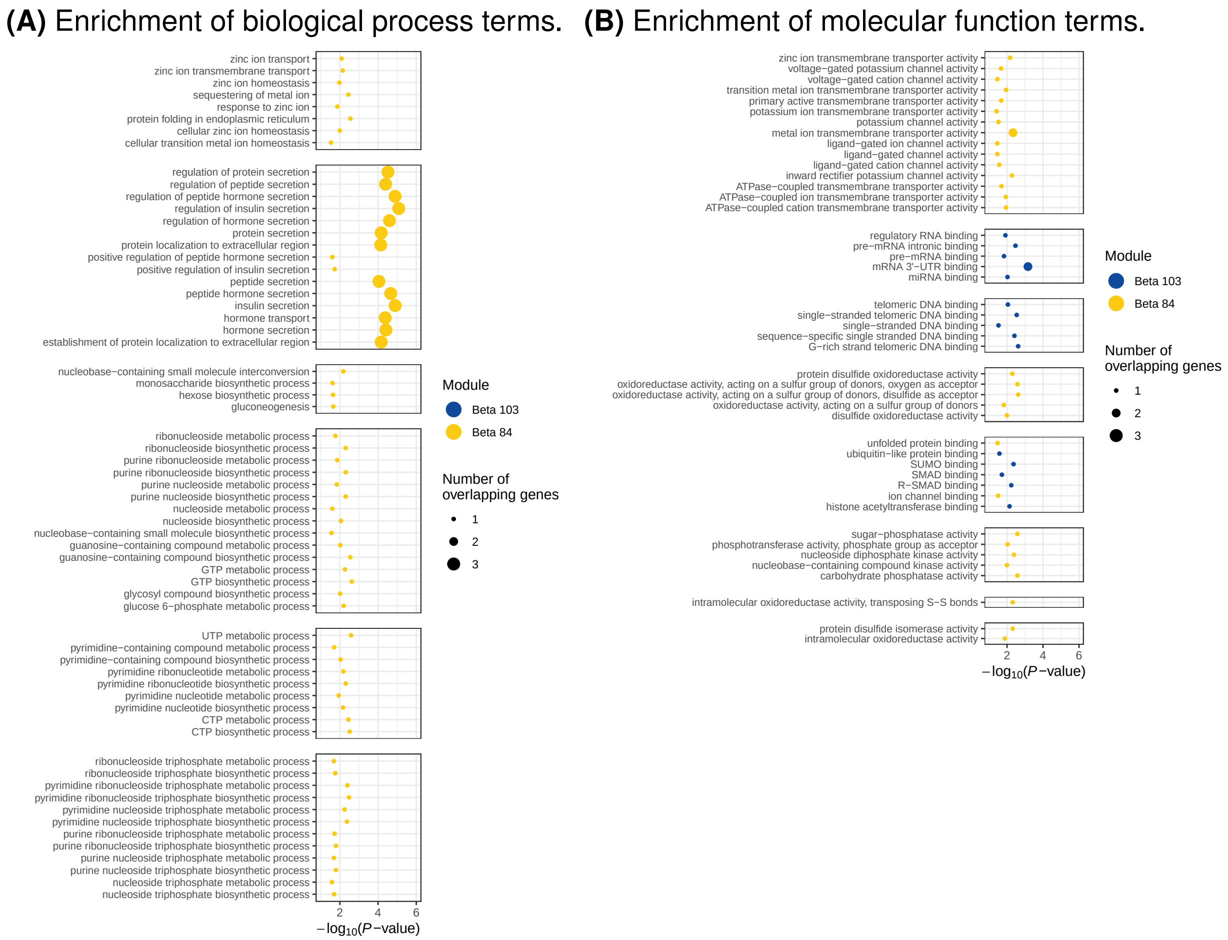


**Fig. S11. Enrichment of gene ontology terms in T2D-related gene expression modules.** Enrichment (x-axis) of gene ontology (GO) terms (FDR<5%; y-axis), faceted by the similarity of genes annotated within each GO term. Point size corresponds to the number of genes within the GO term that overlap genes in an expression module. Color indicates gene expression module. (A) Biological process terms. (B) Molecular function terms.

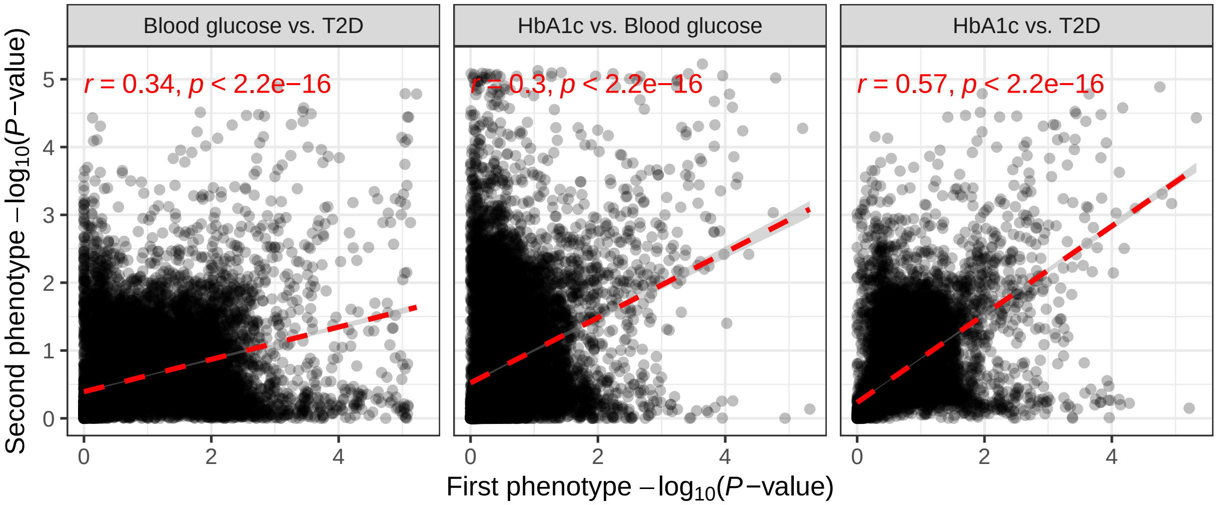


**Fig. S12. Comparison of polygenic priority score results across phenotypes.** Comparison of polygenic priority score empirical -$\log_{10}$(*P*-values) for T2D and T2D-related traits. X-axis corresponds to the first phenotype listed in facets, y-axis corresponds to the second phenotype listed in facets.

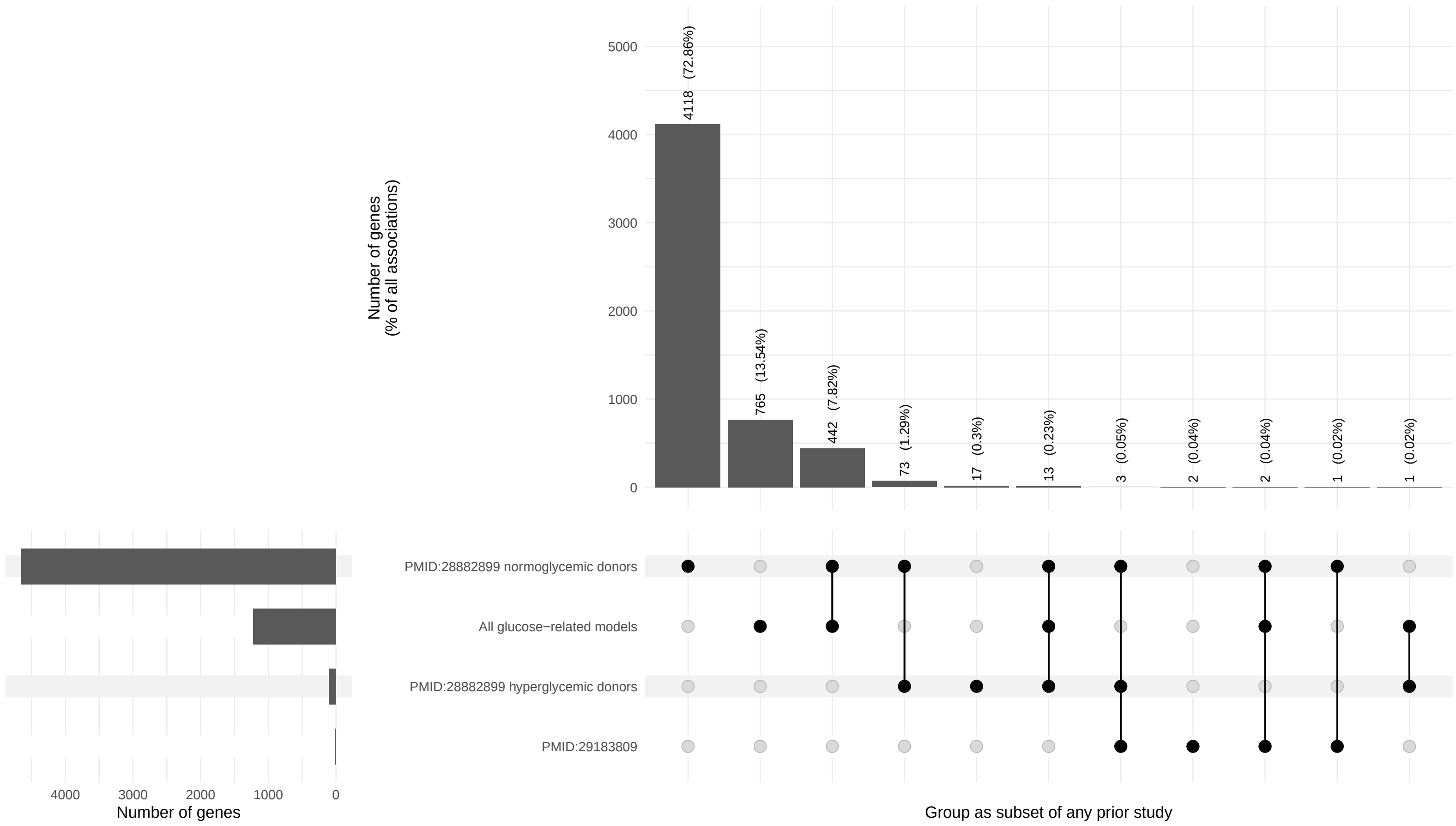


**Fig. S13. Overlap of differential gene expression results across studies.** Number of associated genes (FDR<5%; y-axis) shared between studies (x-axis). Percent of all associations reported within parenthesis of main barplot. "All glucose-related models" refers to all glucose-related models reported in this study across all cell types and time points: LvH, continuous glucose, and continuous time:glucose interaction models.

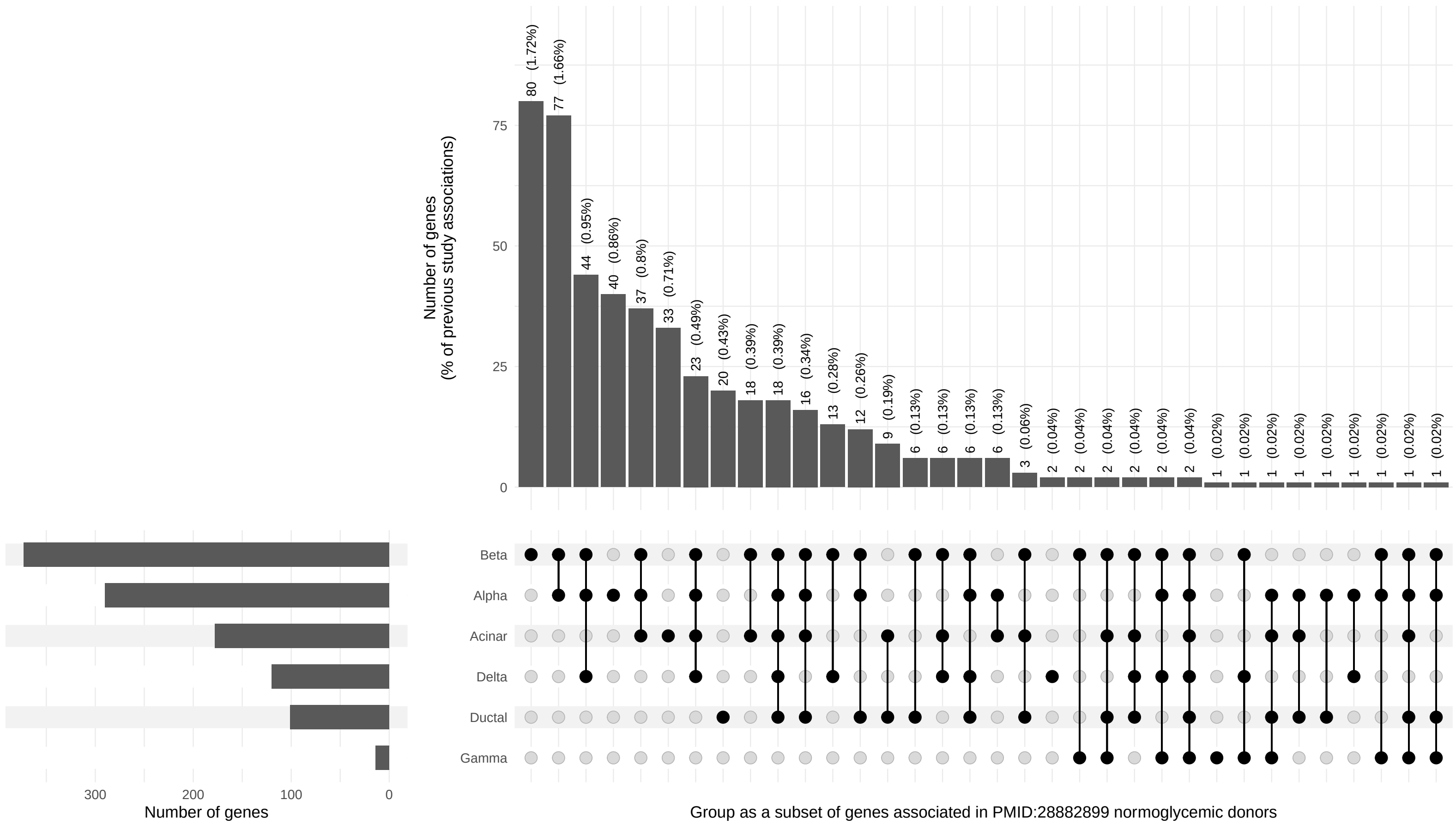


**Fig. S14. Overlap of differential gene expression results with normoglycemic donor results from PMID:28882899.** Number of associated genes (FDR$<$5%; y-axis) from normoglycemic donors in PMID:28882899 identified in glucose-related models from this study (LvH, continuous glucose, and continuous time:glucose interaction models) across cell types (x-axis).

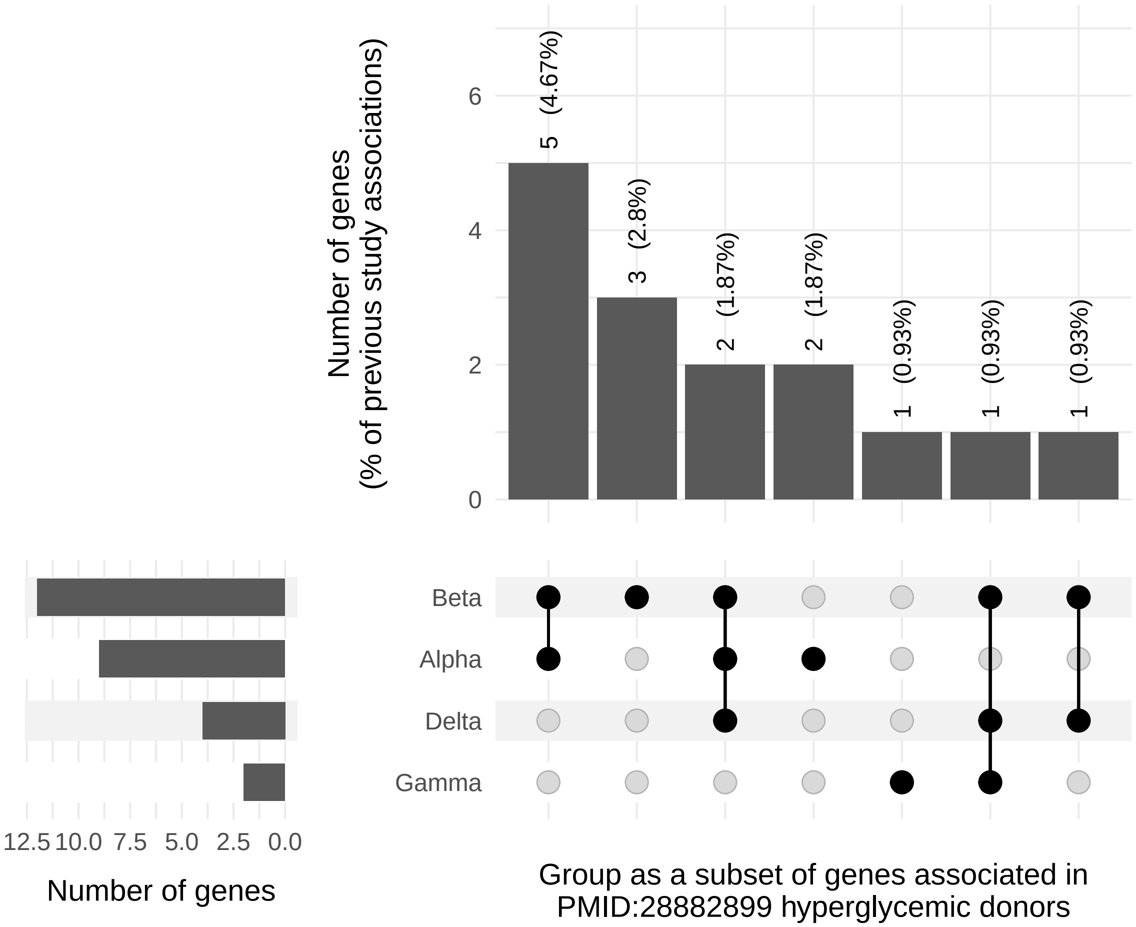


**Fig. S15. Overlap of differential gene expression results with hyperglycemic donor results from PMID:28882899.** Number of associated genes (FDR$<$5%; y-axis) from hyperglycemic donors in PMID:28882899 identified in glucose-related models from this study (LvH, continuous glucose, and continuous time:glucose interaction models) across cell types (x-axis).

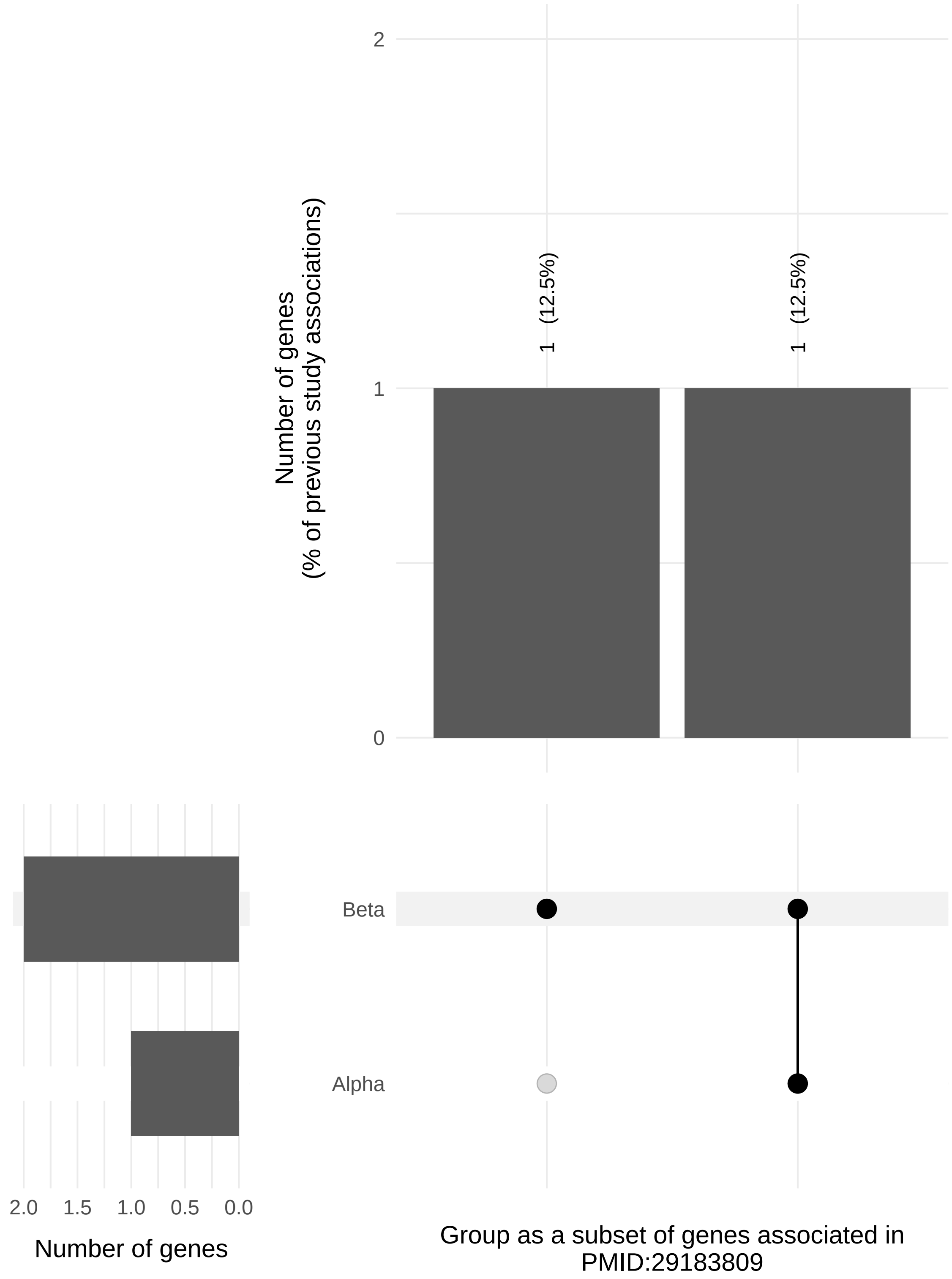


**Fig. S16. Overlap of differential gene expression results with results from PMID:29183809.** Number of associated genes (FDR<5%; y-axis) in PMID:29183809 identified in glucose-related models from this study (LvH, continuous glucose, and continuous time:glucose interaction models) across cell types (x-axis).

| | | |
|---|---|---|
| Donor | HP18227 | HP19208 |
| Age (years) | 36 | 35 |
| Sex | Male | Male |
| Body mass index (kg/m$^2$) | 27.6 | 21.7 |
| HbA1c | 5.8% | 4.7% |
| Source of islets | Prodo Labs | Prodo Labs |
| History of diabetes | No | No |
| Cause of death | Trauma (vehicle accident) | Stroke |
| Islet purity (%) | 85% | 85% |
| Islet viability (%) | 95% | 95% |
| 10x Genomics reagent kit | SC3'v2 | SC3'v3 |

**Table S1. Donor characteristics.** Characteristics of the human pancreatic islet donors for this experiment.

| Cell type | Number of modules | Median module size |
|---|---|---|
| Acinar | 52 | 7 |
| Alpha | 66 | 7 |
| Beta | 58 | 7 |
| Delta | 52 | 6 |
| Ductal | 71 | 8 |
| Gamma | 48 | 6 |

**Table S2. Summary of gene expression modules.** Total number of expression modules identified in each cell type along with median number of genes in the module.
